# Phyllosphere exudates select for distinct microbiome members in sorghum epicuticular wax and aerial root mucilage

**DOI:** 10.1101/2022.07.18.500406

**Authors:** Marco E. Mechan-Llontop, John Mullet, Ashley Shade

## Abstract

Phyllosphere exudates create specialized microhabitats that shape microbial community diversity. Here, we explore the microbiome associated with two sorghum phyllosphere exudates, the epicuticular wax and aerial root mucilage. We hypothesized that these exudates selectively enrich for microbiome members that support host resilience to stress. Thus, we assessed the microbiome associated with the epicuticular wax from sorghum plants under non-limiting and limiting water conditions, and the aerial root mucilage from nitrogen-fertilized and non-fertilized plants. In parallel, we isolated and characterized hundreds of bacteria from wax and mucilage, and integrated data from cultivation-independent and -dependent approaches to gain deeper insights into phyllosphere functions and phenotypes. We found that *Sphingomonadaceae* and *Rhizobiaceae* families were the major taxa in the wax regardless of water availability to plants and that plant development only modestly affected wax bacterial community structure. The mucilage-associated bacterial microbiome contained several described diazotrophic species, and its structure was strongly influenced by sorghum development but only modestly influenced by fertilization. In contrast, the fungal community structure of mucilage was strongly affected by the year of sampling but not by fertilization or plant developmental stage, suggesting a decoupling of fungal-bacterial dynamics in the mucilage. Our bacterial isolate collection from wax and mucilage increased phylogenetic diversity of non-rhizosphere, plant-associated bacteria by ~20% from previous work, and several isolates matched 100% to detected amplicon sequence variants. This work expands our understanding of the microbiome of phyllosphere exudates and advances our long-term goal of translating microbiome research to support sorghum cultivation for biofuel production.

## INTRODUCTION

The phyllosphere, which includes the above-ground plant structures, has diverse surface features (Ruinen 1965; Vacher et al. 2016; Doan et al. 2020). It is a microbial habitat that is exposed to rapid environmental fluctuations and stressors, including UV exposure, temperature and extreme fluctuations in nutrient and water availability. Thus, the diversity and functions of the phyllosphere microbiome reflects this complex habitat (Lindow and Brandl 2003; Vorholt 2012; Vacher et al. 2016). To adapt to abiotic stresses, plants produce a diversity of exudates on their external surfaces (Chai and Schachtman 2022). The secreted exudates vary in composition and structure, creating specialized phyllosphere microhabitats (Galloway et al. 2020). Exudates that accumulate in the phyllosphere include epicuticular wax on stems and leaves (Kunst and Samuels 2003), sugar-rich mucilage on aerial root structures (Bennett et al. 2020), floral nectaries (Rering et al. 2018), and extrafloral nectaries in stems and leaves (Pierce 2019). Because of their potential as locations of microbial engagement with the host, research has been initiated to explore these microbial communities that reside on phyllosphere exudates.

In the current study, we investigate the microbiome of epicuticular wax and aerial root mucilage. Plants secrete epicuticular wax on leaves, leaf sheaths, and stems for prevention of water loss under drought stress (Xue et al. 2017), reflection of solar radiation (Steinmüller and Tevini 1985), and pathogen protection (Serrano et al. 2014; Wang et al. 2020). Epicuticular waxes are enriched in long-chain hydrocarbons. The major wax components include alkanes, alcohols, esters, and fatty acids, as well as varying levels of triterpenoids, sterols, and flavonoids (von Wettstein-Knowles 1974; Kunst and Samuels 2003)(Busta et al., 2020) The wax composition and quantities are affected by plant species, plant developmental stage, and environmental conditions (Yeats and Rose 2013). It has been shown that epicuticular waxes affect bacterial and fungal plant colonization in a species-dependent manner (Beattie and Marcell 2002; Tsuba et al. 2002). Also, wax accumulation and composition directly impact the phyllosphere diversity at the community level (Reisberg et al. 2013). Although epicuticular wax plays multiple roles in plant resilience to abiotic stress, our understanding of its community diversity is limited. A study in *Arabidopsis thaliana* found that Proteobacteria, Bacteroidetes, and Actinobacteria are the dominant phyla associated with wax on leaves. Described wax phyllosphere bacterial families include Flavobacteriaceae, Flexibacteriaceae, Methylobacteriaceae, Rhizobiaceae, Sphingomonadaceae, Enterobacteriaceae, and Pseudomonadaceae (Reisberg et al. 2013). However, how bacterial communities are organized in wax topography and whether bacteria can metabolize epicuticular wax are unknown.

Plants also secrete abundant polysaccharide-rich mucilage on phyllosphere compartments on aerial roots and the above ground portion of brace roots. Brace roots support plant anchorage as well as water and nutrient uptake (Stamp and Kiel 1992; Ku et al. 2012; Reneau et al. 2020). In 2018, van Deynze et al. 2018 reported that the mucilage of aerial roots of a landrace maize harbor a diazotroph microbiota that captures almost 80% of the nitrogen needed by the host from the atmosphere. The bacterial genera *Acinetobacter*, *Agrobacterium, Enterobacter, Klebsiella, Lactococcus, Pantoea, Pseudomonas, Rahnella, Raoultella, Stenotrophomonas*, and others have been found in association with the mucilage of maize. These bacteria are capable of biological nitrogen fixation (BNF), synthesizing indole-3-Acetic Acid (IAA), utilizing 1-amino-1-cyclopropanecarboxylic acid (ACC), and solubilizing phosphates. The unique polysaccharide composition of the mucilage may modulate its associated microbiota (van Deynze et al. 2018; Higdon et al. 2020b). The maize mucilage is enriched in a mixture of monosaccharides including fucose (28%), galactose (22%), arabinose (15%), glucuronic acid (11%), xylose (11%), mannose (8%), glucose (1%) and galacturonic acid (1%) (van Deynze et al. 2018; Amicucci et al. 2019). The polysaccharide composition of root mucilage may vary between maize genotypes and change depending on environmental conditions (Nazari et al. 2020).

Bioenergy sorghum (*Sorghum bicolor* L. Moench) is a heat and drought-tolerant annual crop being developed for production of biomass, biofuels and bioproducts (Mullet et al. 2014; Nelle et al. 2019). Bioenergy sorghum confers 75%-90% greenhouse gas mitigation when used for ethanol production or biopower generation respectively (Olson et al. 2012), but excess nitrogen fertilizer is required to grow it, resulting in the release of nitrous oxide and relatively lower carbon benefit than other biofuel feedstocks that do not have high fertilizer demands (Kent et al. 2020 and Scully et al. 2021). In the 1980s, it was hypothesized that the mucilage secreted by sorghum aerial roots harbors diazotroph bacteria, as has been shown in the landrace maize (Bennett et al. 2020), but evidence of this in sorghum remains limited. Although the polysaccharide composition of the sorghum aerial root mucilage is uncharacterized, it is expected that the sorghum mucilage is similar in composition to maize (van Deynze et al. 2018; Amicucci et al. 2019). Taken together, it is expected that understanding microbiome interactions on the sorghum mucilage may provide insights into microbiome-enabled solutions to optimize diazotrophic nitrogen to the host and, in parallel, reduce nitrogen fertilizer needs for bioenergy sorghum.

During development, bioenergy sorghum accumulates high levels of epicuticular wax on stems and leaves to adapt to biotic and abiotic stress, including resistance to water loss and potential pathogen exclusion. Sorghum epicuticular wax chemistry and structure have been extensively studied. The accumulation, and composition of sorghum epicuticular wax are affected by several factors, including plant age, genotype, water availability, and environmental stresses (Bianchi et al. 1978; Avato et al. 1984; Jordan et al. 1984; Steinmüller and Tevini 1985; Shepherd et al. 1995; Jenks et al. 1996; Bondada et al. 1996; Shepherd and Wynne Griffiths 2006; Xue et al. 2017). However, the influence of wax topography and chemistry on bacteria colonization and community structure is unknown.

In the present study, we investigate the microbiome structure associated with bioenergy sorghum epicuticular wax and aerial root mucilage. We hypothesized that the wax and mucilage select for microbiome members that have traits that support host drought tolerance and nutrient uptake among other functions. To begin examining this hypothesis, the microbiome composition and structure of these specialized exudates was analyzed using field studies and treatments the modulate plant water and nitrogen status. Specifically, we assessed the bacterial microbiome associated with the epicuticular wax from sorghum plants under both non-limiting and limiting water availability, and the bacterial and fungal microbiomes associated with the aerial root mucilage from nitrogen (N)-fertilized and non-fertilized sorghum plants. In addition, we curated a bacterial isolate collection from each phyllosphere exudate to compare diversity and to assess how well standard cultivation conditions cover the cultivation-independent microbiome diversity so that potential targets for microbiome management could be prioritized. We integrate data from both cultivation-independent and -dependent approaches to gain deeper insights into the microbiome diversity and dynamics of sorghum epicuticular wax and aerial root mucilage.

## METHODS

### Collection of sorghum stems and recovery of epicuticular wax

We collected samples from the bioenergy sorghum (*Sorghum bicolor*) hybrid TX08001 grown at the Texas A&M University Research Farm in College Station, Texas (30°55’5.55” N, 96°.43’64.6” W). Sorghum plants were grown in 5 replicate 32 rows by 30 m plots at standard planting density and fertilization (Olson et al., 2012). We sampled replicate plots 1-5 at 60 (08/03/2020) and 90 (09/02/2020) days after plant emergence (DAE). While sorghum plants at 60 DAE were irrigated to maintain non-limiting water status, plants at 90 DAE were grown without irrigation to induce water-limiting conditions until harvesting. We collected stem sections that were covered in epicuticular wax, using razor blades to destructively sample the fifth and sixth fully elongated stem node-internodes below the growing zone into sterile whirl-pak bags. In total, we collected 50 stem samples during the growing season of 2020. All samples were kept on ice for transport, shipped on dry ice to Michigan State University, and then stored at −80 °C. We used sterile razor blades to carefully remove and collect the epicuticular wax from stems in sterile 1.5 ml Eppendorf tubes. Epicuticular wax samples were stored at −80 °C until processing.

### Collection of sorghum aerial roots and removal of the mucilage

We collected samples from the bioenergy sorghum cultivar TAM 17651 grown at the Great Lakes Bioenergy Research Center (GLBRC), as part of the Biofuel Cropping System Experiment (BCSE) in Hickory Corners, Michigan (42°23’41.6” N, 85°22’23.1” W). Sorghum plants were grown in 5 replicate 30×40 m plots arrayed in a randomized complete block design. Within each plot, nitrogen fertilizer-free subplots were maintained either in the western or eastern -most 3m of each plot. We sampled replicate plots 1-4 in both the main and nitrogen-fertilizer free subplots at 60 and 90 DAE. We used sterile razor blades to carefully collect between 3 to 5 aerial nodal roots per plant that were covered with visible mucilage into sterile 50 ml Eppendorf tubes. In total, we collected 180 aerial root samples during the growing seasons of 2020 and 2021. All samples were kept on ice for transport, and then stored at −80 °C. In the laboratory, we added 15 ml of sterile distilled water and kept the roots for 5 min at room temperature to fully hydrate the aerial root mucilage. We collected 1 ml of mucilage into sterile 1.5 ml Eppendorf tubes per sample. Mucilage samples were stored at −80 °C until processing.

### Culturing the epicuticular wax and mucilage microbiomes

For bacterial isolation, we pooled the epicuticular wax collected from different plants, as described above, and resuspended 100 mg of wax in 1 ml of sterile distilled water. We also pooled the mucilage collected from different plants, as described above. To capture a diversity of bacteria, 100 μl of the wax suspension and the mucilage dilutions 10^-1^ to 10^-4^ were separately plated in duplicate on a variety of culture media **(Supplementary Table 2)**. All plates were incubated both at 25°C and 37°C for up to 14 days. To select for anaerobic bacteria, agar plates were placed in anaerobic jars (Mitsubishi AnaeroPack 7.0L rectangular jar) containing three bags of anaerobic gas generator (Thermo Scientific AnaeroPack Anaerobic Gas generator) and incubated at 25°C and 37°C for up to 14 days. To enrich for bacteria resistant to desiccation, one hundred microliters of dilution 10^-1^ from the wax and mucilage were inoculated on 20 ml of 50% TSB liquid culture supplemented with 6000 polyethylene-glycol. To enrich for bacteria resistant to terpenoids, 100 ml of dilution 10^-1^ from the wax and mucilage were inoculated on 20 ml of 50% TSB liquid culture supplemented with 1% (v/v) of either linalool or B-caryophyllene. Liquid cultures were incubated at 28°C for 24 h, and dilutions 10^-1^ to 10^-4^ were plated in duplicate on R2A agar plates for 24 h. Well isolated individual colonies were picked with a sterile toothpick and transferred to a new R2A plate. To confirm bacterial purity, individual bacterial colonies were transferred three times on new R2A agar plates. Glycerol stock (25% v/v) of pure bacteria isolates were stored at −80°C.

### Metagenomic DNA extraction and amplicon sequencing

Microbial DNA was extracted from 0.5 ml of mucilage and 100 mg of epicuticular wax using a DNeasy PowerSoil kit (Qiagen, Maryland, USA) according to the manufacturer’s instructions. To confirm successful DNA extraction, the metagenomic DNA was quantified using a qubit 2.0 fluorometer (Invitrogen, Carlsbad, CA, USA), and visualized in a 1% agarose gel. Then, the PCR amplifications and sequencing of the V4 region of the 16S rRNA bacterial or archaeal gene from the epicuticular wax and mucilage samples and the ITS1 region of the fungal rRNA gene from the mucilage samples only were performed. DNA concentrations were normalized to approximately 1 μg/μl between all samples before PCR amplification and sequencing. The V4 hypervariable region of the 16S rRNA gene was amplified using the universal primers 515F (5’-GTGCCAGCMGCCGCGGTAA-3’) and 806R (5’-GGACTACHVGGGTWTCTAAT-3’) (Gregory et al. 2011)under the following conditions: 95°C for 3 min, followed by 30 cycles of 95°C for 45 s, 50°C for 60 s, and 72°C for 90 s, with a final extension at 72°C for 10 min. The metagenomic DNA of each sample was submitted to the Genomics Core of the Research Technology Support Facility at Michigan State University for library preparation and sequencing using the Illumina MiSeq platform v2 Standard flow cell in a 2×250bp paired-end format, using their standard operating protocol.

The ITS1 region was amplified using primers ITS1f (5’-CTTGGTCATTTAGAGGAAGTAA□3’) and ITS2 (5’-GCTGCGTTCTTCATCGATGC-3’) (Smith and Peay 2014) with the addition of index adapters CS1-TS-F: 5’ – ACACTGACGACATGGTTCTACA – [TS-For] – 3’ and CS2-TS-R: 5’ – TACGGTAGCAGAGACTTGGTCT – [TS-Rev] – 3’ as requested by the Genomics Sequencing Core under the following PCR conditions: 94°C for 3 min, followed by 35 cycles of 94°C for 30 s, 52°C for 30 s, and 68°C for 30 s, with a final extension at 68°C for 10 min. The amplification was performed with GoTaq Green Master Mix (Promega). The PCR products were purified with ExoSAP-IT reagent, and sample sequencing was completed by the Genomics Core of the Research Technology Support Facility at Michigan State University using the Illumina MiSeq platform v2 Standard flow cell in a 2×250bp paired-end format. For quality control purposes, positive and negative controls were included throughout the DNA extraction, PCR amplification, and sequencing processes. A 75 μl aliquot of the ZymoBIOMICS Microbial Community Standard (Zymo Research, Irvine, CA, U.S.A) and 75 μl aliquot of an in-house Community Standard (Colovas et al. 2022) were included as positive controls. Sterile DEPC-treated water was included as negative control.

### Bacterial genomic DNA extraction

Bacteria colonies that were first streaked and isolated for purity were grown on 2 ml of 50% TSB liquid culture at 28°C for 24 h. Bacteria culture was centrifuged at 5,000 rpm for 10 min. Genomic DNA of each isolate was extracted by using the Zymo – Quick DNA Fungal/Bacterial 96 kit following the manufacturer’s protocol. Total genomic DNA was quantified using a qubit 2.0 fluorometer and visualized in a 1% agarose gel. The PCR amplification of the full-length 16S rRNA gene with universal primers 27F (5’-AGAGTTTGATCCTGGCTCAG-3’) and 1492R (5’-TACGGTTACCTTGTTACGACTT-3’) (Miller et al. 2013)was performed by using the Pfu Turbo DNA polymerase (Agilent) under the following conditions: 95°C for 2 min, followed by 24 cycles of 95°C for 30 s, 48°C for 30 s, and 72°C for 3 min, with a final extension at 72°C for 10 min. PCR products were purified with ExoSAP-IT reagent and submitted for Sanger sequencing at the Genomics Core of the Research Technology Support Facility at Michigan State University, MI, USA.

### Bacterial and fungal amplicon sequencing analysis

Paired-end sequencing data from each sequencing experiment were processed with Qiime2 (Bolyen et al. 2019) version 2021.8.0. In brief, sequences were imported using the PairedEndFastqManifestPhred33V2 format. Sequence quality control, denoising, and generation of feature tables containing counts for the Amplicon Sequencing Variants (ASVs) were performed with the q2-dada2 plugin version 2021.8.0 (Callahan et al. 2016). Trimming parameters for the DADA2 plugin were selected with FIGARO version 1.1.2 (Weinstein et al. 2019). ASVs tables and representative sequences from each sequencing experiment were merged with the q2-feature-table plugin. ASV taxonomy (of merged ASVs) was assigned with the q2-feature-classifier plugin using the SILVA version 1.38 database (Quast et al. 2013) for bacteria and UNITE version 8.3 database (Nilsson et al. 2019) for fungi.

ASVs table, taxonomy table, and sample metadata files were imported into R version 4.1.3 for data visualization and statistical analysis. Diversity and statistical analyses were performed using the phyloseq (McMurdie and Holmes 2013) and vegan (Dixon 2003) packages. Differential abundance analysis was performed with the DESeq2 package (Love et al. 2014).

### Full-length 16S rRNA gene Sanger sequencing analysis: Culturing phyllosphere exudate microbiota

To generate a consensus sequence of the full-length 16S rRNA gene from each bacterial isolate, sequences were imported into Geneious version 2021.2.2 (https://www.geneious.com/). High-quality forward and reverse sequences were aligned and trimmed to generate a consensus sequence. Then, the consensus sequence was searched with BLAST for taxonomic classification. Cd-hit version 4.8.1 (Li and Godzik 2006) was used to remove redundant 16S rRNA sequences. To identify bacterial isolates that match 100% to the identified ASVs from the culture-independent approach, a local BLAST search was performed. In summary, a local BLAST database was created with all non-redundant 16S rRNA sequences from our bacterial collection using the *makeblastdb* command and the *-dbtype nucl* option. A BLAST search was carried out to identify related sequences in the representative sequences (ASVs dna-sequences.fasta) file generated from the DADA2 denoising step with the *blastn* command, and the flowing options: “6 qseqid sseqid pident length mismatch gapopen qstart qend sstart send evalue bitscore”.

### Comparison with publicly available plant-associated bacterial genomes

We retrieved 637 plant-associated (PA) bacterial genomes that were classified as non-root associated from the (Levy et al. 2017) study. High quality bacterial genomes were annotated with Prokka (Seemann 2014) using an in-house python script and annotated 16S rRNA gene copies were identified. For bacteria with multiple 16S rRNA copies, Cd-hit version 4.8.1 (Li and Godzik 2006) was used to remove redundant sequences (99% similarity) and one 16S rRNA sequence was conserved, totaling 433 unique PA sequences. All 16S rRNA sequences from the PA bacterial genome dataset were concatenated in a single fasta file with the *cat* command. Cd-hit was used to remove redundant sequences (100 % similarity) from the 16S rRNA concatenated file. All non-redundant 16S rRNA sequences from both the sorghum bacterial collections and the publicly available PA bacteria were merged in a single *fasta* file. Sequence alignment was performed with MAFFT v7.407 (Katoh et al. 2002). Alignment trimming was performed with trimAl (Capella-Gutiérrez et al. 2009). A maximum-likelihood (ML)-based phylogenetic tree was built with IQ-TREE 2.2.0-beta version (Minh et al. 2020). ModelFinder version (-m TEST option) (Kalyaanamoorthy et al. 2017) was used to select the best model for the phylogenetic tree construction. Branch support was assessed using 1,000 ultrafast boostrap approximations (-bb 1000 option) (Hoang et al. 2018). Phylogenetic diversities were calculated as the total tree length, that represents the expected number of substitutions per site. Phylogenetic tree was edited with iTOLs version 6.5.8 (Letunic and Bork 2021).

### Data and code availability

The data analysis workflows for sequence processing and ecological statistics are available on GitHub (https://github.com/ShadeLab/Sorghum_phyllosphere_microbiome_MechanLlontop_2022.git). Raw sequencing data has been deposited in the Sequence Read Archive NCBI database under BioProject accession number PRJNA844896 (including 16S rRNA and ITS amplicons). Full-length 16S rRNA sequence data has been deposited in the Genbank with accession numbers ON973084-ON973283.

## RESULTS

### Sequencing summary

In total, we sequenced the bacterial 16S rRNA V4 region from 48 epicuticular wax samples from the 2020 growing season, as well as the bacterial 16S rRNA V4 region from 179 mucilage samples and the fungal ITS region from 173 mucilage samples that were collected across two growing seasons in 2020 and 2021. We obtained 8,648,839 bacterial sequences from the wax, and 20,606,039 bacterial and 20,181,404 fungal sequences from the mucilage. After quality control, removal of chimeras, and denoising, 7,930,768 quality bacterial reads were obtained from the wax samples, and 19,880,634 bacterial and 12,157,819 fungal sequences were obtained from mucilage **(Table 1)**. For wax, the total number of sequences per sample after the denoising process with DADA2 into Amplicon Sequence Variants (ASVs) ranged from 1,722 to 272,108. After the removal of nonbacterial and unassigned sequences, a total of 2,386,033 sequences remained, with sequencing reads per wax sample ranging from 138 to 206,128. We removed wax samples with fewer than 1000 sequences, and the remaining 42 epicuticular wax samples were rarefied to 1,303 sequences for further analysis **(Figure 1A)**.

**Figure 1.**
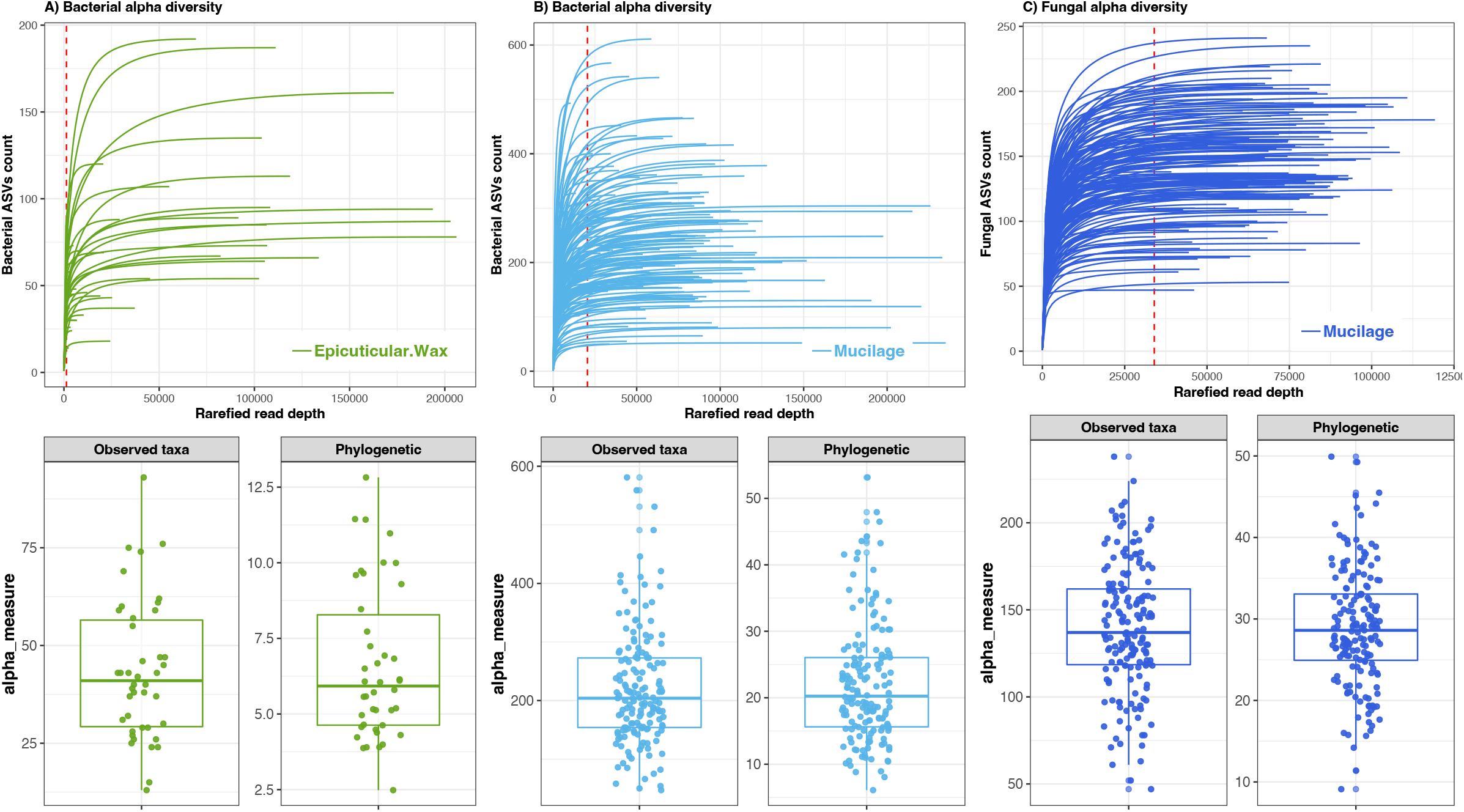
Sequencing effort and alpha diversity for sorghum epicuticular wax and aerial root mucilage. Top: Rarefaction curves of quality-controlled sequences. Bottom: *Observed taxa* and *Phylogenetic diversity* (PD) metrics. A) Epicuticular wax samples were rarefied to 1,303 reads per sample (red vertical line). Amplicon sequencing variants (ASVs) were defined at 100% identity of 16S rRNA gene. B) Aerial root mucilage samples were rarefied to 20,519 reads per sample (red vertical line). Amplicon sequencing variants (ASVs) were defined at 100% identity of 16S rRNA gene. C) Aerial root mucilage samples were rarefied to 33,975 reads per sample (red vertical line). Amplicon sequencing variants (ASVs) were defined at 100% identity of ITS1 gene.

**Table 1.** Sequencing summary of sorghum wax and mucilage microbial communities characterized in this study.

|  | Epicuticular<br>wax (16S<br>rRNA) | Aerial root<br>mucilage (16S<br>rRNA) | Aerial root<br>mucilage (16S<br>rRNA) | Aerial root<br>mucilage<br>(ITS1) | Aerial root<br>mucilage<br>(ITS1) |
| --- | --- | --- | --- | --- | --- |
|  | 2020 | 2020 | 2021 | 2020 | 2021 |
| Number of samples | 48 | 99 | 80 | 92 | 81 |
| Raw Read Pairs | 8,648,839 | 12,783,054 | 10,034,885 | 10,403,184 | 9,778,220 |
| QC reads<br>% | 7,930,768 | 10,809,135 | 9,071,499 | 6,200,571 | 5,957,248 |
| Chloroplast/Mitochondria/<br>unassigned of QC reads | 70% | 24% | 48% | 0% | 0% |

For roots mucilage, the number of bacterial sequences per sample after the denoising ranged from 222 to 330,853. After the removal of nonbacterial and unassigned sequences, a total of 12,956,774 sequences remained, with sequencing reads per sample ranging from 110 to 235,069. We removed samples with fewer than 20,000 sequences, and the remaining 158 samples were rarefied to 20,519 sequences for comparative analysis **(Figure 1B)**. The number of fungal sequences per mucilage sample after the denoising ranged from 78 to 119,207. After the removal of non-fungal and unassigned sequences, a total of 12,297,453 sequences remained, with sequencing reads per sample ranging from 32 to 119,207. We filtered mucilage samples with fewer than 30,000 ITS sequences, and the remaining 171 samples were rarefied to 33,975 sequences for comparative analysis **(Figure 1C)**.

### *Sphingomonadaceae* and *Rhizobiaceae* dominate sorghum epicuticular wax bacterial diversity

Altogether, we identified 534 bacterial ASVs in epicuticular wax. Wax bacterial microbiome samples collected from sorghum plants at 60 DAE and 90 DAE had different richness (observed taxa p= 0.03) and Chao1 indices (p= 0.03), but not Shannon diversity (P_Shannon_ = 0.48, comparisons using Wilcoxon rank) **(Supplementary Table 1)**. There was higher variation in the community structure in the epicuticular wax on plants at 90 DAE compared with plants at 60 DAE (betadispers F=17.921, p=0.001). There was a small but significant influence of sorghum developmental stage on the epicuticular wax community structure (R2=0.064, p-value= 0.003, PERMANOVA **Figure 2A**, **Table 2)**.

**Figure 2.**
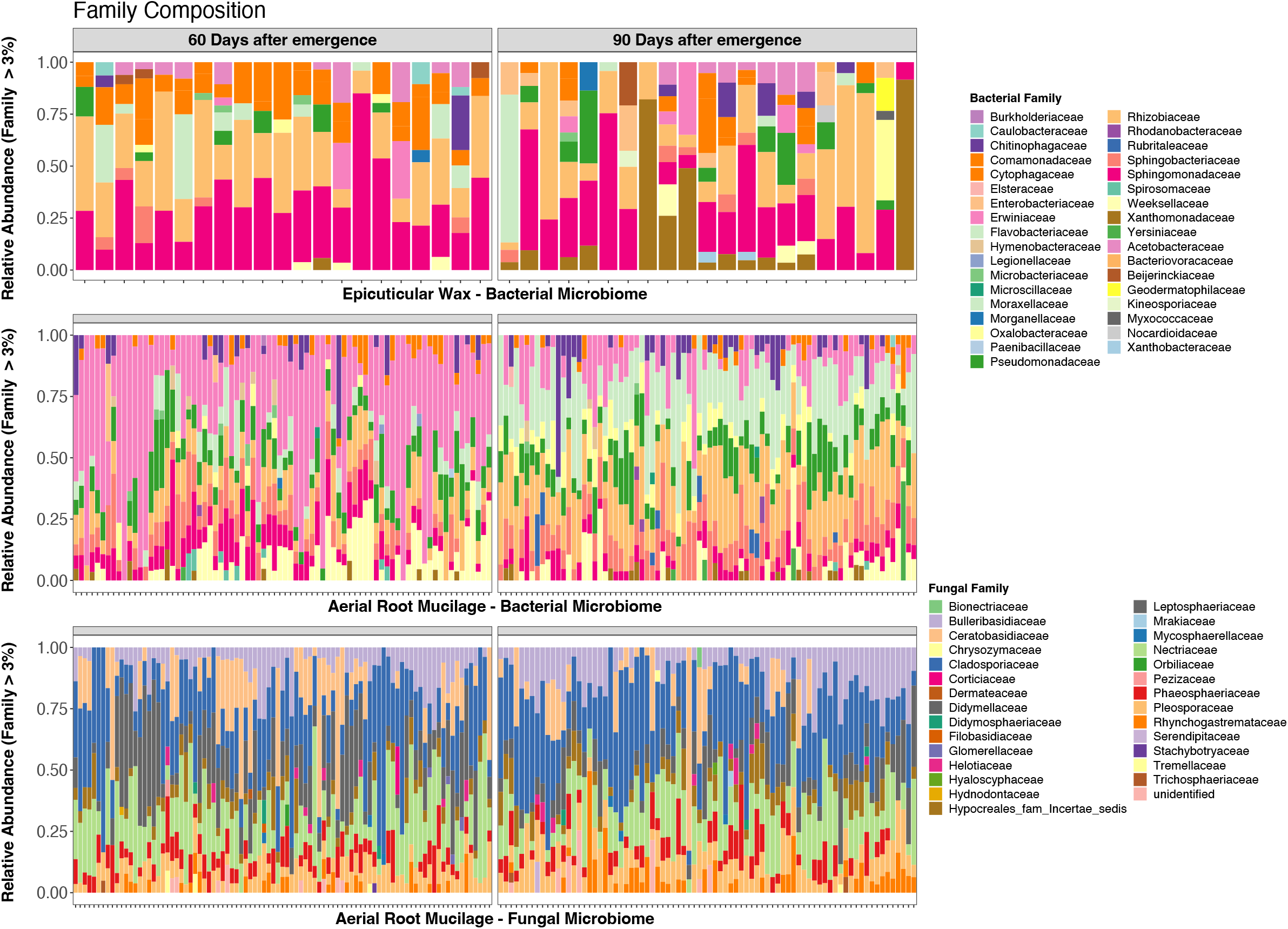
Relative abundance (RA) of microbial taxa in sorghum epicuticular wax and aerial root mucilage. A) RA of Epicuticular wax bacterial communities. Samples were rarefied to 1,303 reads per sample. B) RA of aerial root mucilage bacterial communities. Samples were rarefied to 20,519 reads per sample. C) RA of aerial root mucilage fungal communities. Samples were rarefied to 33,975 reads per sample. Only families with RA > 3% are shown.

**Table 2.**
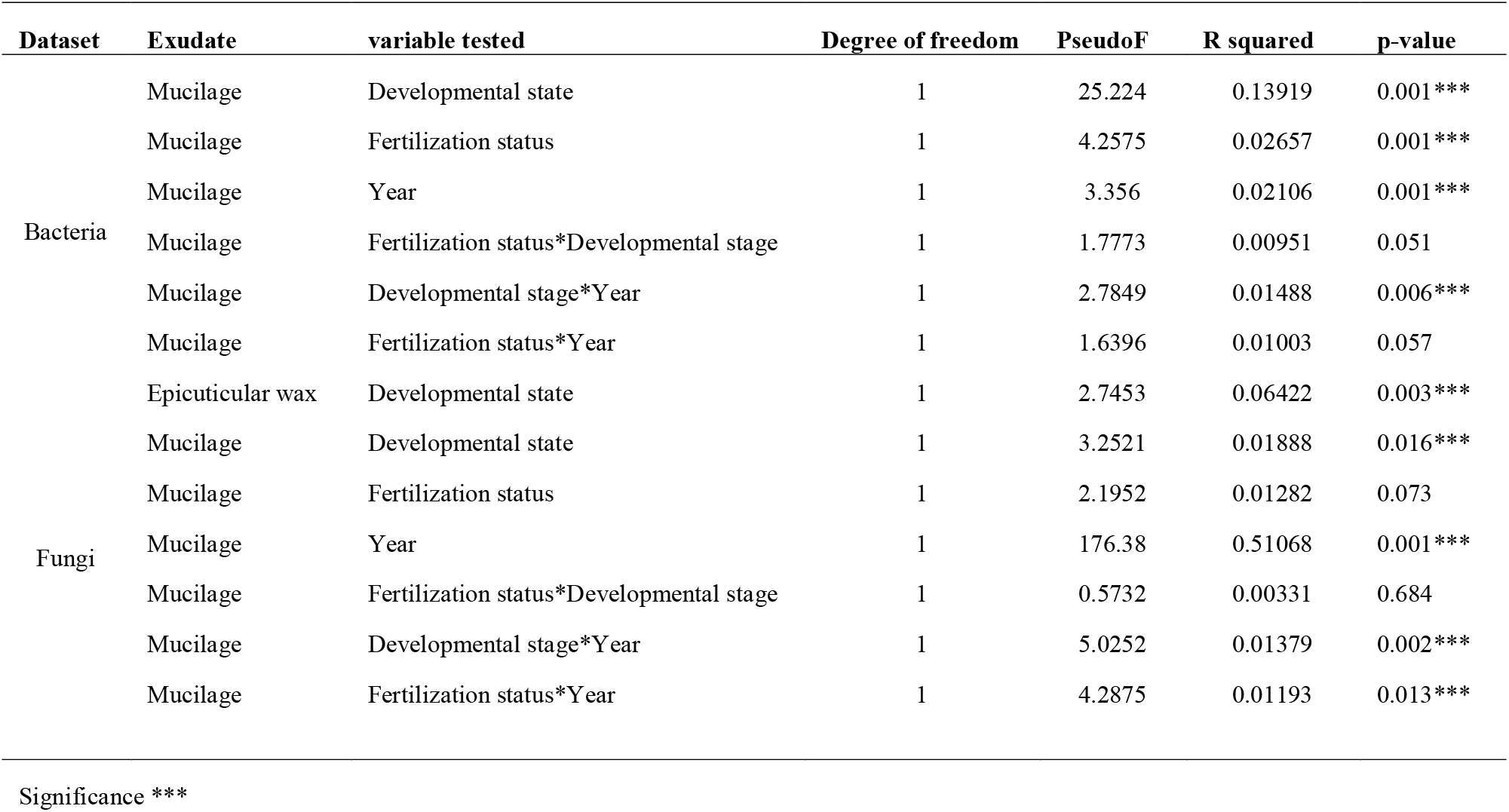
Permuted multivariate analysis of variance (PERMANOVA) results for hypothesis tests for differences in beta diversity.

The sorghum epicuticular wax microbiome was dominated by the Proteobacteria (84% mean relative abundance) and Bacteroidetes (11%) bacteria phyla. The bacterial classes Alphaproteobacteria (54%), Gammaproteobacteria (30%), and Bacteroidia (11%) were in highest abundance. Sphingomonadaceae (25%), Rhizobiaceae (21%), and Xanthomonadaceae (7%) were the major bacterial families in sorghum epicuticular wax **(Figure 3A).** At the genus level, *Sphingomonas* (28%), *Rhizobium* (12%), *Aureimonas* (10%), and *Acinetobacter* (5%) were the dominant taxa in wax **(Supplementary Figure 1)**. Differential abundance analysis with DESeq showed that only 1 ASV from the Microbacteriacea family was significantly more abundant on the wax of plants at 60 DAE, and that 1 ASV assigned to *Pseudoxanthomonas* genera was significantly more abundant on the wax of plants at 90 DAE (data not shown).

**Figure 3.**
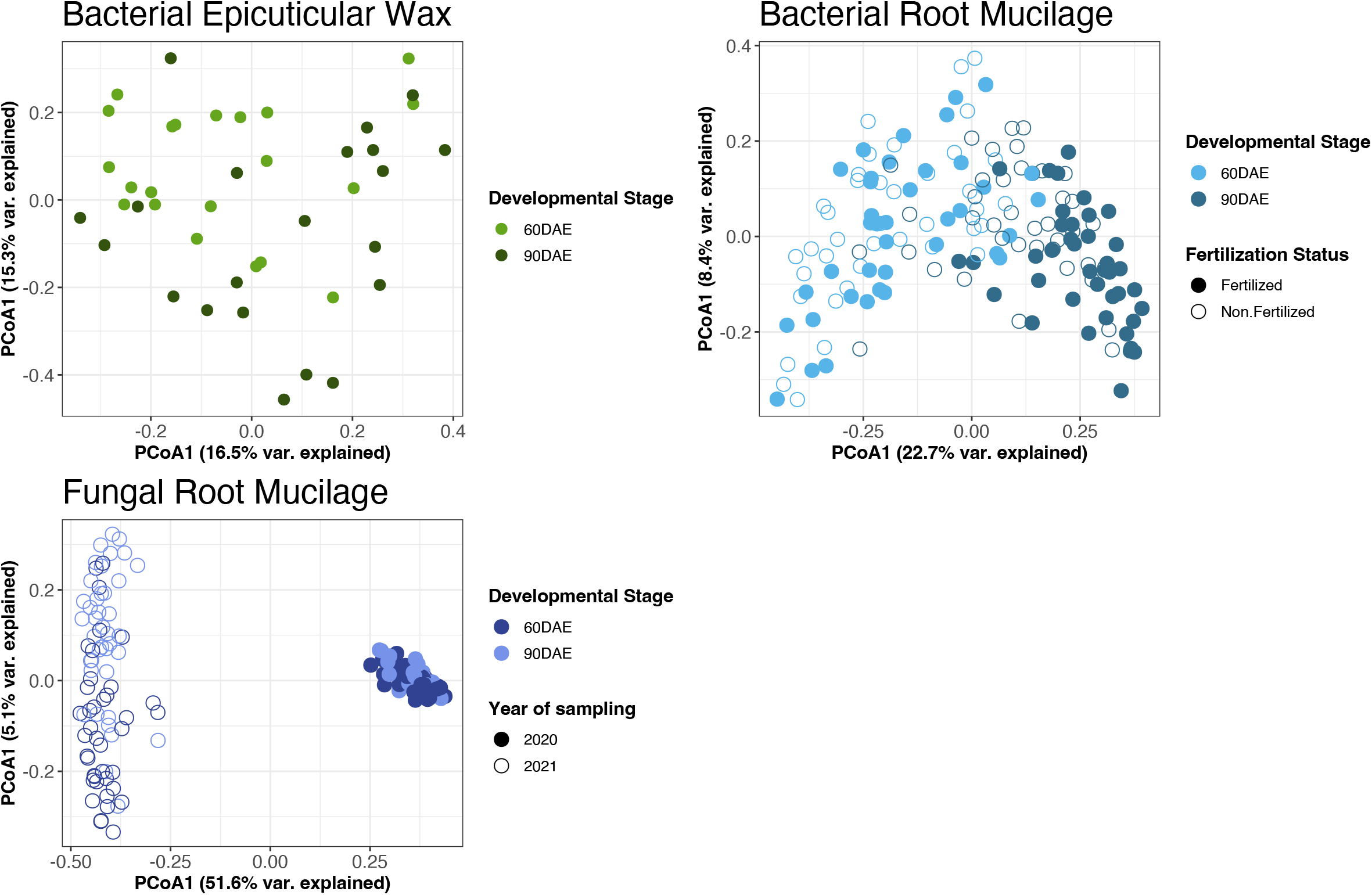
Beta diversity metrics based on the Bray-Curtis distance via Principal Coordinates Analysis (PCoA) for sorghum epicuticular wax and aerial root mucilage. A) PCoA of the epicuticular wax bacterial communities, B) PCoA of the aerial root mucilage bacterial communities, C) PCoA of the aerial root mucilage fungal communities. A multivariate permutational ANOVA test (PERMANOVA) using Adonis from Bray Curtis dissimilarity was performed for each data set **(Table 2).**

### The sorghum aerial root mucilage includes diazotroph-associated bacterial species

Altogether, 12,047 bacterial ASVs were identified in aerial root mucilage. There were differences in Shannon diversity between mucilage samples collected from sorghum plants at 60 DAE and 90 DAE (P_Shannon_= 0.00002, comparisons using Wilcoxon rank), but not for richness (observed species, P = 0.82) or Chao1 (P= 0.48). In contrast, no difference was found between mucilage samples from nitrogen-fertilized plants compared with unfertilized plants. **(Supplementary Table 1)**. There was different dispersion in community structure (beta dispersion) by plant developmental stage (F=19.56, p=0.001) but not by fertilization status (F=1.827, p=0.187). The mucilage bacterial microbiome structure was better explained by developmental stage than fertilization status (adonis R2= 0.13935 v. 0.0267, respectively, both p-value= 0.001) **(Figure 2B).**

The aerial root mucilage bacterial microbiome was dominated by the Proteobacteria (61% mean relative abundance) and Bacteroidota (36%) bacteria phyla. The bacterial class Gammaproteobacteria (40%), Bacteroidia (34%), and Alphaproteobacterial (21%) were the most abundant. Erwiniaceae (23%), Rhizobiaceae (14%), Flavobacteriaceae (12%), Pseudomonadaceae (9%), and Sphingomonadaceae (6%) were the major bacterial families in mucilage. There were several genera associated with diazotroph activity detected, including *Acinetobacter, Curtobacterium, Herbaspirillum, Pantoea, Pseudomonas, Strenotrophomonas*, and others **(Supplementary Figure 1).** A differential abundance analysis with the DESeq package (*p-value* = 0.01) identified 25 ASVs enriched on the mucilage of plants at 60 DAE and 72 ASVs significantly enriched in plants at 90 DAE **(Figure 4)**.

**Figure 4.**
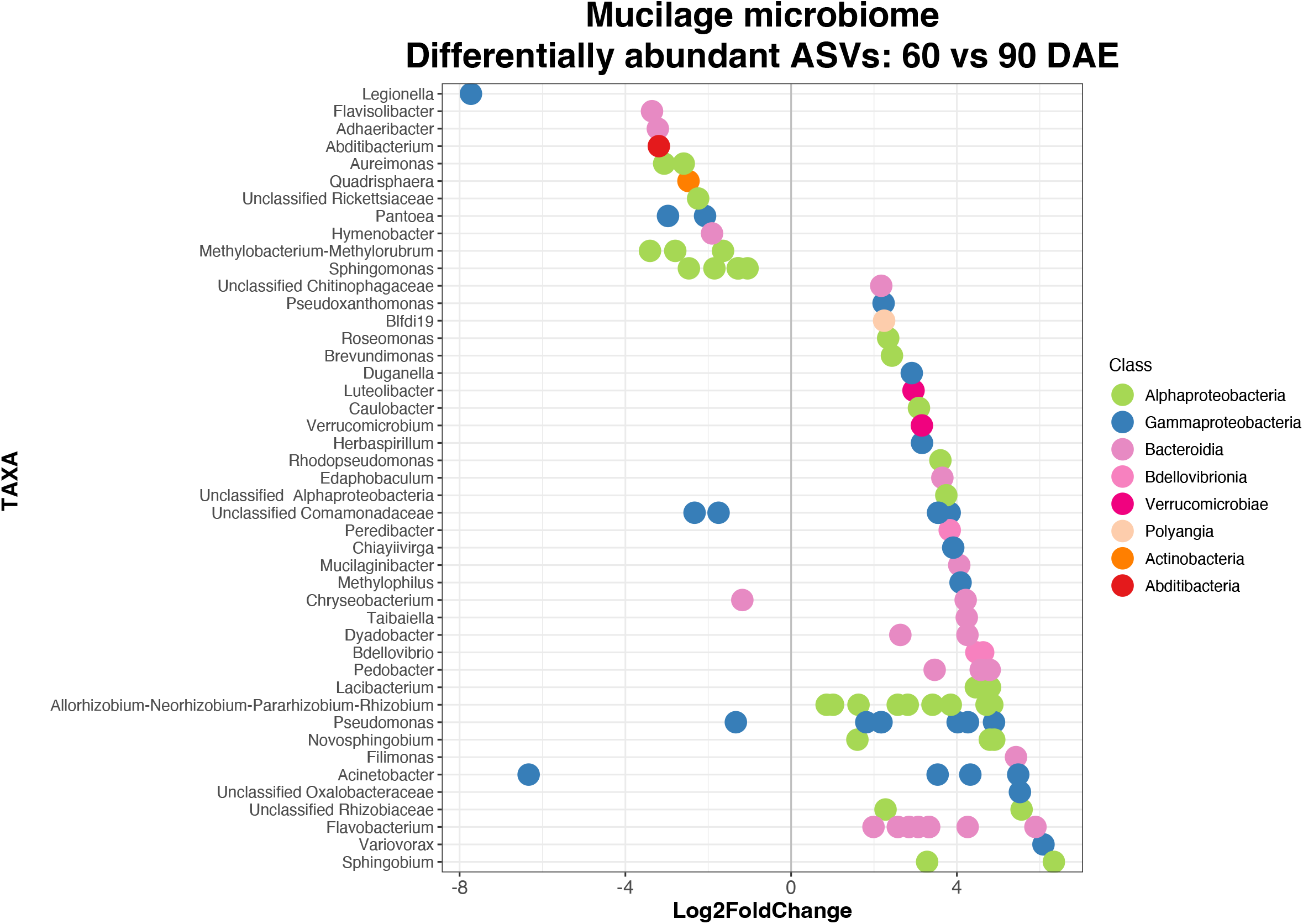
Differential abundance analysis at the level of Amplicon sequencing variants (ASVs). Differentially enriched ASVs in the aerial root mucilage of plants at 60 and 90 DAE are shown. The fold change is shown on the x-axis and bacterial taxa are listed on the y-axis. Each colored dot represents a separate ASV.

### Fungal diversity in the sorghum aerial root mucilage

Altogether, 5,641 fungal ASVs were identified in aerial root mucilage. There were differences in richness and diversity between mucilage samples collected from sorghum plants during the 2020 and 2021 growing seasons (P_Observed_ = 0.008, P_Shannon_ = 0.004). Differences were also in richness between mucilage samples from nitrogen-fertilized plants compared with unfertilized plants (P_Chao_ = 0.004, P_Observed_ = 0.006), but not for Shannon diversity. No difference was observed between mucilage samples from plants at 60 DAE vs. 90 DAE **(Supplementary Table 1)**. The mucilage fungal microbiome structure was strongly influenced by year of collection (adonis R2= 0.51068, p-value= 0.001). Fungal community structure was slightly influenced by developmental stage (R2= 0.01888, p-value= 0.015), but not by fertilization status (R2= 0.01282, p-value= 0.081) **(Figure 2C).**

The mucilage fungal microbiome was dominated by the Ascomycota (76%) and Basidiomycota (23.7%) phyla. The Dothideomycetes (50%), Sordariomycetes (24%), and Tremellomycetes (14%) fungal classes were the most abundant. Cladosporium (22%), Nectriaceae (17%), Didymellaceae (14%), Bulleribasidiaceae (9 %), Pleosporaceae (8%) were the dominant fungal families in the mucilage. The genera *Cladosporium* exhibited higher abundance in the 2020 growing season (34%) compared with 2021 (14%). In contrast, we found an enrichment of the genera *Epicoccum* in 2021 (18%) compared with the 2020 growing season (0.02%) **(Supplementary Figure 1)**.

### Sorghum phyllosphere wax and mucilage harbor unique bacterial diversity

Bacterial culture collections from the epicuticular wax and aerial root mucilage were constructed by enriching bacteria with putative plant-beneficial traits **(Supplementary Figure 2)**. In total, 500 bacteria from the wax and 800 bacteria from the mucilage were isolated, and then a subset of 200 isolates from both the wax and mucilage were taxonomically identified by sequencing the full-length 16S rRNA gene. The wax bacterial collection was dominated by the Proteobacteria, followed by Actinobacteria, and Bacteroidetes phyla, and the mucilage bacterial collection was dominated by the Proteobacteria, followed by Actinobacteria, Firmicutes, and Bacteroidetes phyla **(Supplementary Table 4)**.

The bacterial diversity found in the culture-independent approach (ASV table) was compared with the diversity found in the wax and mucilage bacterial collections. Forty-eight ASVs matched with 100% identity to strains in the isolate collections **(Table 3)**. Most of the bacterial families found in the sorghum wax and mucilage had representatives among the isolate collection **(Figure 5)**. While some lineages, such as the Microbacteriaceae family, were overrepresented in the wax bacterial collection, others like the Rhizobiaceae and Sphingomonadaceae families were underrepresented. Families such as Beijerincklaceae, Chitinophagaceae, Oxalobacteraceae, and others were not captured by our initial cultivation efforts. Similarly, while the Microbacteriaceae and Enterobacteriaceae families were overrepresented in the mucilage bacterial collection, the Flavobacteriaceae, and Sphingomonadaceae families were underrepresented. Families such as Cytophagaceae, and Oxalobacteraceae were not captured by our initial cultivation efforts.

**Figure 5.**
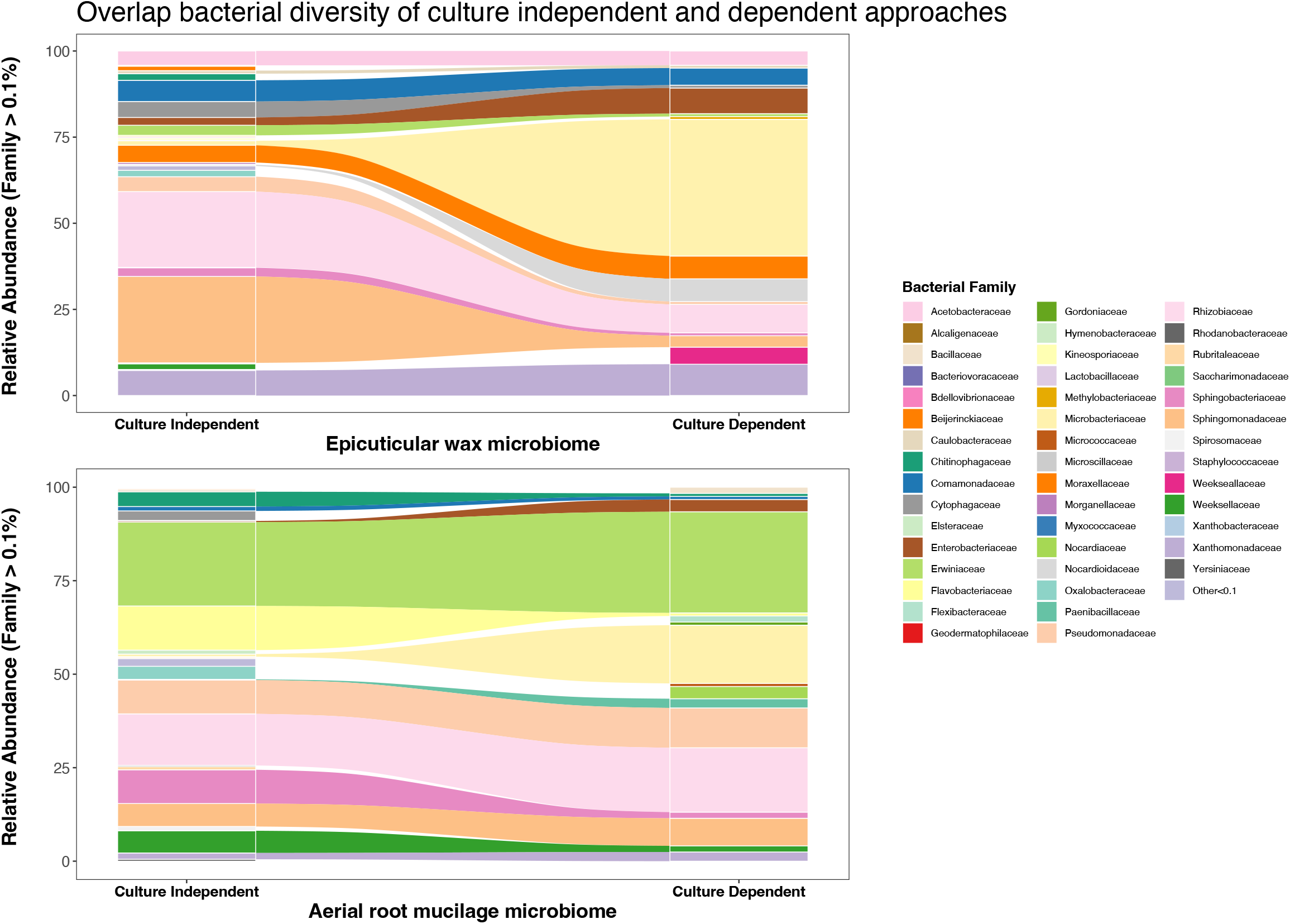
Overlap bacterial diversity found in the sorghum epicuticular wax and aerial root mucilage based on culture-independent and culture-dependent approaches. Relative abundance at the family level > 0.1% are shown.

**Table 3.** List of bacterial ASVs (Amplicon Sequencing Variants) with 100% similarity to the 16S rRNA gene from the bacterial isolates.

| qseqid | sseqid | pide<br>nt | lengt<br>h | mis-<br>matc<br>h | gapop<br>en | qsta<br>rt | qen<br>d | ssta<br>rt | sen<br>d | evalue | bitsco<br>re |
| --- | --- | --- | --- | --- | --- | --- | --- | --- | --- | --- | --- |
| 04ecfad5772d2e09a84a0f5ef460536c | SORGH_AS983_conse<br>nsus | 100 | 253 | 0 | 0 | 1 | 253 | 346 | 598 | 2.96E-134 | 468 |
| 09de3dde6c465b74bcb8e6f8a478e4e0 | SORGH_AS768_conse<br>nsus | 100 | 253 | 0 | 0 | 1 | 253 | 320 | 572 | 2.96E-134 | 468 |
| 09de3dde6c465b74bcb8e6f8a478e4e0 | SORGH_AS810_conse<br>nsus | 100 | 253 | 0 | 0 | 1 | 253 | 412 | 664 | 2.96E-134 | 468 |
| 0f18144d308ada95632ab5193d92073f | SORGH_AS191_conse<br>nsus | 100 | 253 | 0 | 0 | 1 | 253 | 453 | 705 | 2.96E-134 | 468 |
| 10afda2baef44de4c584a6641de399b1 | SORGH_AS789_conse<br>nsus | 100 | 253 | 0 | 0 | 1 | 253 | 405 | 657 | 2.96E-134 | 468 |
| 10afda2baef44de4c584a6641de399b1 | SORGH_AS950_conse<br>nsus | 100 | 253 | 0 | 0 | 1 | 253 | 406 | 658 | 2.96E-134 | 468 |
| 10afda2baef44de4c584a6641de399b1 | SORGH_AS879_conse<br>nsus | 100 | 253 | 0 | 0 | 1 | 253 | 364 | 616 | 2.96E-134 | 468 |
| 10afda2baef44de4c584a6641de399b1 | SORGH_AS965_conse<br>nsus | 100 | 253 | 0 | 0 | 1 | 253 | 363 | 615 | 2.96E-134 | 468 |
| 119b70614392764602bb20f8bbe8f960 | SORGH_AS659_conse<br>nsus | 100 | 253 | 0 | 0 | 1 | 253 | 449 | 701 | 2.96E-134 | 468 |
| 1bc599cac5647c5c30496e93ca551557 | SORGH_AS189_conse<br>nsus | 100 | 253 | 0 | 0 | 1 | 253 | 433 | 685 | 2.96E-134 | 468 |
| 1bc599cac5647c5c30496e93ca551557 | SORGH_AS194_conse<br>nsus | 100 | 253 | 0 | 0 | 1 | 253 | 271 | 523 | 2.96E-134 | 468 |
| 1fa19a198819b145e52bb266e4e7840b | SORGH_AS613_conse<br>nsus | 100 | 253 | 0 | 0 | 1 | 253 | 381 | 633 | 2.96E-134 | 468 |
| 26c35a7f1d51ff8a47651ff85e8614fe | SORGH_AS565_conse<br>nsus | 100 | 253 | 0 | 0 | 1 | 253 | 401 | 653 | 2.96E-134 | 468 |
| 2d77cf80c9b03b82ce7ed6eacfa04e12 | SORGH_AS909_conse<br>nsus | 100 | 253 | 0 | 0 | 1 | 253 | 449 | 701 | 2.96E-134 | 468 |
| 392807149dd6b2ddc5a33b220b32ed27 | SORGH_AS1015_cons<br>ensus | 100 | 253 | 0 | 0 | 1 | 253 | 335 | 587 | 2.96E-134 | 468 |
| 428e4a80a1e14a59d72810a32f62f434 | SORGH_AS199_conse<br>nsus | 100 | 253 | 0 | 0 | 1 | 253 | 472 | 724 | 2.96E-134 | 468 |
| 48454bd504b23d75d289a23f12faa874 | SORGH_AS211_conse<br>nsus | 100 | 253 | 0 | 0 | 1 | 253 | 468 | 720 | 2.96E-134 | 468 |
| 496ecde24f9ab698992413d3d4f04b5f | SORGH_AS457_conse<br>nsus | 100 | 253 | 0 | 0 | 1 | 253 | 364 | 616 | 2.96E-134 | 468 |
| 496ecde24f9ab698992413d3d4f04b5f | SORGH_AS282_conse<br>nsus | 100 | 253 | 0 | 0 | 1 | 253 | 452 | 704 | 2.96E-134 | 468 |
| 49d46d00a93443b060707ab2db8ba82d | SORGH_AS936_conse<br>nsus | 100 | 253 | 0 | 0 | 1 | 253 | 366 | 618 | 2.96E-134 | 468 |
| 4e0481f68111cd3463386f70247922bb | SORGH_AS447_conse<br>nsus | 100 | 253 | 0 | 0 | 1 | 253 | 436 | 688 | 2.96E-134 | 468 |
| 4e0481f68111cd3463386f70247922bb | SORGH_AS1171_cons<br>ensus | 100 | 253 | 0 | 0 | 1 | 253 | 385 | 637 | 2.96E-134 | 468 |
| 5673ae66819e3bdbbc09461e4959159e | SORGH_AS997_conse<br>nsus | 100 | 253 | 0 | 0 | 1 | 253 | 478 | 730 | 2.96E-134 | 468 |
| 5673ae66819e3bdbbc09461e4959159e | SORGH_AS1094_cons<br>ensus | 100 | 253 | 0 | 0 | 1 | 253 | 451 | 703 | 2.96E-134 | 468 |
| 6c5f4f9666d82b1bedf94e99acfe6741 | SORGH_AS510_conse<br>nsus | 100 | 253 | 0 | 0 | 1 | 253 | 441 | 693 | 2.96E-134 | 468 |
| 73373d2d94551b3c53c3e01b6a02aeb3 | SORGH_AS859_conse<br>nsus | 100 | 253 | 0 | 0 | 1 | 253 | 415 | 667 | 2.96E-134 | 468 |
| 73373d2d94551b3c53c3e01b6a02aeb3 | SORGH_AS923_conse<br>nsus | 100 | 253 | 0 | 0 | 1 | 253 | 396 | 648 | 2.96E-134 | 468 |
| 73373d2d94551b3c53c3e01b6a02aeb3 | SORGH_AS1208_cons<br>ensus | 100 | 253 | 0 | 0 | 1 | 253 | 386 | 638 | 2.96E-134 | 468 |
| 73373d2d94551b3c53c3e01b6a02aeb3 | SORGH_AS888_conse<br>nsus | 100 | 253 | 0 | 0 | 1 | 253 | 412 | 664 | 2.96E-134 | 468 |
| 73373d2d94551b3c53c3e01b6a02aeb3 | SORGH_AS209_conse<br>nsus | 100 | 253 | 0 | 0 | 1 | 253 | 308 | 560 | 2.96E-134 | 468 |
| 7521edd61e578d0d49cfd01fbf8cc2f5 | SORGH_AS860_conse<br>nsus | 100 | 253 | 0 | 0 | 1 | 253 | 468 | 720 | 2.96E-134 | 468 |
| 7521edd61e578d0d49cfd01fbf8cc2f5 | SORGH_AS852_conse<br>nsus | 100 | 253 | 0 | 0 | 1 | 253 | 423 | 675 | 2.96E-134 | 468 |
| 777c50185200d35116740a777e462069 | SORGH_AS887_conse nsus | 100 | 253 | 0 | 0 | 1 | 253 | 419 | 671 | 2.96E-134 | 468 |
| 777c50185200d35116740a777e462069 | SORGH_AS893_conse nsus | 100 | 253 | 0 | 0 | 1 | 253 | 416 | 668 | 2.96E-134 | 468 |
| 7ff346973a282aa55de296afdb5d74af | SORGH_AS212_conse nsus | 100 | 253 | 0 | 0 | 1 | 253 | 447 | 699 | 2.96E-134 | 468 |
| 8452a6541e06768ed3c1aabf4e6568a7 | SORGH_AS622_conse nsus | 100 | 253 | 0 | 0 | 1 | 253 | 441 | 693 | 2.96E-134 | 468 |
| 8ce12e88f6b59bb09494567f0d678092 | SORGH_AS260_conse nsus | 100 | 253 | 0 | 0 | 1 | 253 | 399 | 651 | 2.96E-134 | 468 |
| 8f820a46cfecd19477f4485d1c436764 | SORGH_AS956_conse nsus | 100 | 253 | 0 | 0 | 1 | 253 | 468 | 720 | 2.96E-134 | 468 |
| 8f820a46cfecd19477f4485d1c436764 | SORGH_AS833_conse nsus | 100 | 253 | 0 | 0 | 1 | 253 | 467 | 719 | 2.96E-134 | 468 |
| 945184b6386c192c0066e0a98a154780 | SORGH_AS313_conse nsus | 100 | 253 | 0 | 0 | 1 | 253 | 466 | 718 | 2.96E-134 | 468 |
| 945184b6386c192c0066e0a98a154780 | SORGH_AS906_conse nsus | 100 | 253 | 0 | 0 | 1 | 253 | 403 | 655 | 2.96E-134 | 468 |
| 945184b6386c192c0066e0a98a154780 | SORGH_AS1146_cons ensus | 100 | 253 | 0 | 0 | 1 | 253 | 384 | 636 | 2.96E-134 | 468 |
| 945184b6386c192c0066e0a98a154780 | SORGH_AS878_conse nsus | 100 | 253 | 0 | 0 | 1 | 253 | 394 | 646 | 2.96E-134 | 468 |
| 945184b6386c192c0066e0a98a154780 | SORGH_AS842_conse nsus | 100 | 253 | 0 | 0 | 1 | 253 | 409 | 661 | 2.96E-134 | 468 |
| 945184b6386c192c0066e0a98a154780 | SORGH_AS1173_cons ensus | 100 | 253 | 0 | 0 | 1 | 253 | 394 | 646 | 2.96E-134 | 468 |
| 945184b6386c192c0066e0a98a154780 | SORGH_AS826_conse nsus | 100 | 253 | 0 | 0 | 1 | 253 | 301 | 553 | 2.96E-134 | 468 |
| 945184b6386c192c0066e0a98a154780 | SORGH_AS834_conse nsus | 100 | 253 | 0 | 0 | 1 | 253 | 391 | 643 | 2.96E-134 | 468 |
| 945184b6386c192c0066e0a98a154780 | SORGH_AS1025_cons ensus | 100 | 253 | 0 | 0 | 1 | 253 | 420 | 672 | 2.96E-134 | 468 |
| 945184b6386c192c0066e0a98a154780 | SORGH_AS1070_cons ensus | 100 | 253 | 0 | 0 | 1 | 253 | 413 | 665 | 2.96E-134 | 468 |
| 9cca219ac7e1d9722ab63545dc219ecd | SORGH_AS835_conse nsus | 100 | 253 | 0 | 0 | 1 | 253 | 437 | 689 | 2.96E-134 | 468 |
| 9cca219ac7e1d9722ab63545dc219ecd | SORGH_AS201_conse nsus | 100 | 253 | 0 | 0 | 1 | 253 | 438 | 690 | 2.96E-134 | 468 |
| 9fe4c3dfc721f87da8ff9661f9600667 | SORGH_AS445_conse nsus | 100 | 253 | 0 | 0 | 1 | 253 | 377 | 629 | 2.96E-134 | 468 |
| a0901407705992dc213ac509ede97d47 | SORGH_AS304_conse nsus | 100 | 253 | 0 | 0 | 1 | 253 | 449 | 701 | 2.96E-134 | 468 |
| a531dc57776d2c54e4dd31f15d2e3517 | SORGH_AS422_conse nsus | 100 | 253 | 0 | 0 | 1 | 253 | 294 | 546 | 2.96E-134 | 468 |
| a537d8bab85c83b0e74c73c55790324b | SORGH_AS993_conse nsus | 100 | 253 | 0 | 0 | 1 | 253 | 407 | 659 | 2.96E-134 | 468 |
| a76e065bdfde0aa883c898ccf5f092fc | SORGH_AS739_conse nsus | 100 | 253 | 0 | 0 | 1 | 253 | 401 | 653 | 2.96E-134 | 468 |
| bdf8a26094624622d68509a87fa75ba7 | SORGH_AS407_conse nsus | 100 | 253 | 0 | 0 | 1 | 253 | 450 | 702 | 2.96E-134 | 468 |
| c2a787cf867ebfdce216002bcc8a94e1 | SORGH_AS442_conse nsus | 100 | 253 | 0 | 0 | 1 | 253 | 193 | 445 | 2.96E-134 | 468 |
| cff60bed359051dd65fe850fd0bd07e1 | SORGH_AS301_00012 | 100 | 253 | 0 | 0 | 1 | 253 | 503 | 755 | 2.96E-134 | 468 |
| d11ef81cd8ab0dfa7a44a92acae1ccfb | SORGH_AS1067_cons ensus | 100 | 253 | 0 | 0 | 1 | 253 | 395 | 647 | 2.96E-134 | 468 |
| d11ef81cd8ab0dfa7a44a92acae1ccfb | SORGH_AS913_conse nsus | 100 | 253 | 0 | 0 | 1 | 253 | 388 | 640 | 2.96E-134 | 468 |
| d11ef81cd8ab0dfa7a44a92acae1ccfb | SORGH_AS1205_cons ensus | 100 | 253 | 0 | 0 | 1 | 253 | 389 | 641 | 2.96E-134 | 468 |
| d11ef81cd8ab0dfa7a44a92acae1ccfb | SORGH_AS864_conse nsus | 100 | 253 | 0 | 0 | 1 | 253 | 407 | 659 | 2.96E-134 | 468 |
| d15bd1dcb9de71d7452648673a59f5b8 | SORGH_AS585_conse nsus | 100 | 253 | 0 | 0 | 1 | 253 | 276 | 528 | 2.96E-134 | 468 |
| d4bcbfee3890222f785195430f9f9dd5 | SORGH_AS321_conse nsus | 100 | 253 | 0 | 0 | 1 | 253 | 218 | 470 | 2.96E-134 | 468 |
| d66072bb4e903a9c1a95728d29427410 | SORGH_AS776_conse nsus | 100 | 253 | 0 | 0 | 1 | 253 | 390 | 642 | 2.96E-134 | 468 |
| d966c575f302342975ba8061cbf50a2c | SORGH_AS853_conse nsus | 100 | 253 | 0 | 0 | 1 | 253 | 419 | 671 | 2.96E-134 | 468 |
| d966c575f302342975ba8061cbf50a2c | SORGH_AS846_conse nsus | 100 | 253 | 0 | 0 | 1 | 253 | 407 | 659 | 2.96E-134 | 468 |
| e1c7d97fe13e9127d225d76a1feb8c78 | SORGH_AS1126_cons ensus | 100 | 253 | 0 | 0 | 1 | 253 | 394 | 646 | 2.96E-134 | 468 |
| e1c7d97fe13e9127d225d76a1fe<br>b8c78 | SORGH_AS787_conse<br>nsus | 100 | 253 | 0 | 0 | 1 | 253 | 389 | 641 | 2.96E-134 | 468 |
| e1c7d97fe13e9127d225d76a1fe<br>b8c78 | SORGH_AS968_conse<br>nsus | 100 | 253 | 0 | 0 | 1 | 253 | 193 | 445 | 2.96E-134 | 468 |
| e34cede80a0f0564aee002c3482f<br>c268 | SORGH_AS1145_cons<br>ensus | 100 | 253 | 0 | 0 | 1 | 253 | 441 | 693 | 2.96E-134 | 468 |
| e34cede80a0f0564aee002c3482f<br>c268 | SORGH_AS961_conse<br>nsus | 100 | 253 | 0 | 0 | 1 | 253 | 412 | 664 | 2.96E-134 | 468 |
| e4c616e0e34cf5f3837203933c1<br>8e498 | SORGH_AS1204_cons<br>ensus | 100 | 253 | 0 | 0 | 1 | 253 | 432 | 684 | 2.96E-134 | 468 |
| e4c616e0e34cf5f3837203933c1<br>8e498 | SORGH_AS421_conse<br>nsus | 100 | 253 | 0 | 0 | 1 | 253 | 432 | 684 | 2.96E-134 | 468 |
| e4c616e0e34cf5f3837203933c1<br>8e498 | SORGH_AS872_conse<br>nsus | 100 | 253 | 0 | 0 | 1 | 253 | 399 | 651 | 2.96E-134 | 468 |
| e4c616e0e34cf5f3837203933c1<br>8e498 | SORGH_AS502_conse<br>nsus | 100 | 253 | 0 | 0 | 1 | 253 | 415 | 667 | 2.96E-134 | 468 |
| e4c616e0e34cf5f3837203933c1<br>8e498 | SORGH_AS875_conse<br>nsus | 100 | 253 | 0 | 0 | 1 | 253 | 378 | 630 | 2.96E-134 | 468 |
| e4c616e0e34cf5f3837203933c1<br>8e498 | SORGH_AS594_conse<br>nsus | 100 | 253 | 0 | 0 | 1 | 253 | 396 | 648 | 2.96E-134 | 468 |
| e4c616e0e34cf5f3837203933c1<br>8e498 | SORGH_AS1077_cons<br>ensus | 100 | 253 | 0 | 0 | 1 | 253 | 424 | 676 | 2.96E-134 | 468 |
| e4c616e0e34cf5f3837203933c1<br>8e498 | SORGH_AS1169_cons<br>ensus | 100 | 253 | 0 | 0 | 1 | 253 | 432 | 684 | 2.96E-134 | 468 |
| e4c616e0e34cf5f3837203933c1<br>8e498 | SORGH_AS612_conse<br>nsus | 100 | 253 | 0 | 0 | 1 | 253 | 425 | 677 | 2.96E-134 | 468 |
| e4c616e0e34cf5f3837203933c1<br>8e498 | SORGH_AS1046_cons<br>ensus | 100 | 253 | 0 | 0 | 1 | 253 | 373 | 625 | 2.96E-134 | 468 |
| e4c616e0e34cf5f3837203933c1<br>8e498 | SORGH_AS1139_cons<br>ensus | 100 | 253 | 0 | 0 | 1 | 253 | 396 | 648 | 2.96E-134 | 468 |
| e4c616e0e34cf5f3837203933c1<br>8e498 | SORGH_AS984_conse<br>nsus | 100 | 253 | 0 | 0 | 1 | 253 | 402 | 654 | 2.96E-134 | 468 |
| e5e0736aa5d9832f64deb09f464<br>2d92f | SORGH_AS1115_cons<br>ensus | 100 | 253 | 0 | 0 | 1 | 253 | 364 | 616 | 2.96E-134 | 468 |
| e7379f7318de2d38ce7c591b86e<br>0a819 | SORGH_AS988_conse<br>nsus | 100 | 253 | 0 | 0 | 1 | 253 | 397 | 649 | 2.96E-134 | 468 |
| e7379f7318de2d38ce7c591b86e<br>0a819 | SORGH_AS930_conse<br>nsus | 100 | 253 | 0 | 0 | 1 | 253 | 438 | 690 | 2.96E-134 | 468 |
| ecedae16c5190f358d2b40c47a9<br>43fa0 | SORGH_AS441_conse<br>nsus | 100 | 253 | 0 | 0 | 1 | 253 | 325 | 577 | 2.96E-134 | 468 |
| f36c50279a8fa1c121cb9dda1ab<br>1cb42 | SORGH_AS335_conse<br>nsus | 100 | 253 | 0 | 0 | 1 | 253 | 417 | 669 | 2.96E-134 | 468 |
| fc6622c636a5210293fb2873fc4<br>761d9 | SORGH_AS745_conse<br>nsus | 100 | 253 | 0 | 0 | 1 | 253 | 393 | 645 | 2.96E-134 | 468 |
| 4c692f4f8bb38b0c0107ff3045fc<br>f123 | SORGH_AS983_conse<br>nsus | 100 | 251 | 0 | 0 | 1 | 251 | 346 | 596 | 3.83E-133 | 464 |

To understand potential novelty and redundancy represented by the diversity of our wax and mucilage bacterial collections, we compared the full-length 16S rRNA genes with those extracted from the bacterial genomes of previously described non-root-associated, plant-associated (PA) bacteria (levy et al. 2017), assigned as non-root-associated. 637 bacterial genomes were retrieved from a publicly available database (see Methods) to provide a reference of context for our 200 sorghum phyllosphere isolates. The final data set contained 527 non-redundant 16S rRNA sequences: 94 new 16S rRNA genes from our wax and mucilage collections, and 433 rRNA genes from the published plant-associated bacterial genomes (**Figure 6**). Our isolates expanded the diversity of described lineages in the collection of plant-associated genomes, as well as sequences that added clades of lineages that were not previously represented in the collection of plant-associated bacterial genomes.

**Figure 6.**
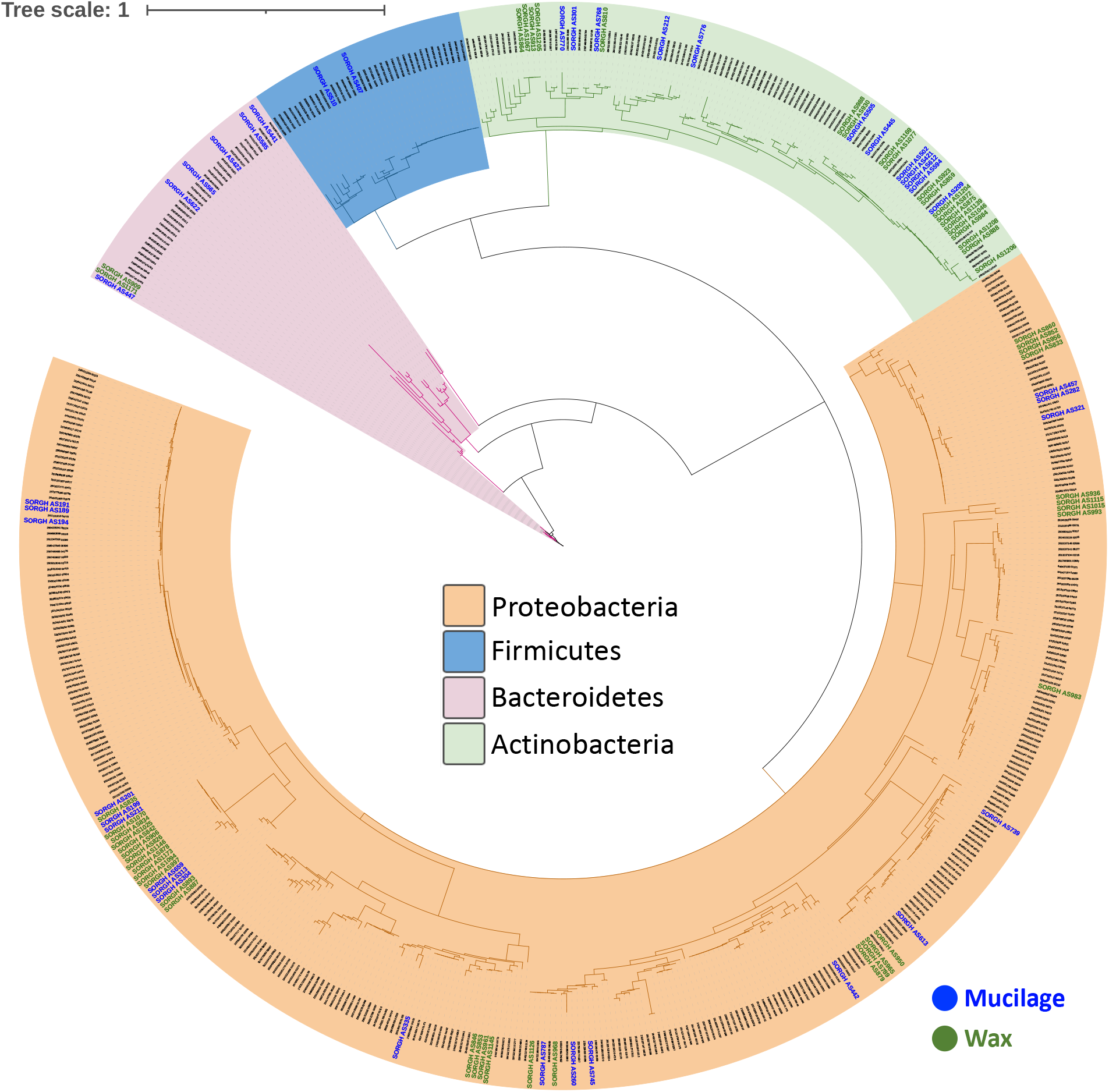
Phylogenetic diversity in the sorghum epicuticular wax and aerial root mucilage. Maximum Likelihood phylogenetic tree (IQTREE, under UNREST+FO+I+G4 model) is based on the 16S rRNA gene alignment.

## DISCUSSION

While our understanding of the interplay between plant exudates and microbiomes has been focused on the rhizosphere compartment, little attention has been paid to the microbiome structure and dynamics in the phyllosphere. To investigate the microbiota associated with phyllosphere exudates, we chose bioenergy sorghum because of its elevated levels of epicuticular wax on stems and leaves and the secretion of mucilage on its aerial roots. In this study, we assessed the microbial communities associated with the epicuticular wax mucilage to test the hypothesis that these exudates select specific microbiome members that support sorghum resilience to stress.

The chemistry of epicuticular wax that covers sorghum stems has been extensively characterized (Bianchi et al. 1978; Jordan et al. 1984; Jenks et al. 2000; Farber et al. 2019a, 2019b), but there is still much to learn about its microbial residents and their colonization dynamics. Thus, we decided to characterize the wax microbiota from stems of field-grown bioenergy sorghum plants at 60 DAE and 90 DAE. We chose these two-time points because they represent different developmental stages. During the vegetative stage, sorghum plants at 60 DAE have all leaves developed and fully expanded. At 90 DAE in the upper mid-west, plants have transition to the reproductive stage, seed development is in progress and nutrients are rapidly relocated to the kernel. The major lineages we detected in the epicuticular stem wax, including *Proteobacteria, Bacteroidetes*, and *Actinobacteria*, agree generally with reports from *Arabidopsis thaliana* and *Sorghum bicolor* epicuticular leaf wax (Reisberg et al. 2013b; Sun et al. 2021). Furthermore, we also observed changes in the relative abundances of several taxa at 60 DAE compared with plants at 90 DAE, which could be associated with changes in the composition of the epicuticular wax as the plant grows (Avato et al. 1984; Jenks et al. 1996), though more work is needed to characterize changes in the chemical composition of the wax alongside the structural changes in the microbiome to understand their relationship more fully. We hypothesize that microbes in wax may be able to metabolize wax components and use them as a carbon source (Ueda et al. 2015), produce hydrocarbons that influence wax chemical composition (Jones 1969; Li et al. 2018), and resist the antimicrobial effect of wax (Shanker et al. 2007). It is possible that wax bacteria, including *Sphingomonas*, *Rhizobium*, and *Pseudomonas* species play role in plant drought resistance as shown in root compartments (Luo et al. 2019; Igiehon et al. 2019; Mahmoudi et al. 2019; Hone et al. 2021). To gain further insight into epicuticular wax microbiome assembly and dynamics, next steps could expand this research not only by including samples from different growing seasons, but also by including sorghum genotypes that are mutants in wax production (Jenks et al. 1994, 2000b; Peters et al. 2009; Punnuri et al. 2017).

For decades it has been suggested that the sorghum aerial root mucilage harbors diazotroph bacteria (Wani 1986; Bennett et al. 2020), but evidence remains limited. We hypothesized that fertilization and developmental stage would strongly influence the phyllosphere mucilage microbiota due to changes in exogenous nutrient availability and changes in mucilage polysaccharide composition. However, our data suggest that that differences in nitrogen fertilization had no notable influence on the microbiome structure for both bacterial and fungal communities. In contrast, plant developmental stage strongly affected the mucilage bacterial microbiome structure. Similar results have been found in other phyllosphere microbiome studies (Copeland et al. 2015; Grady et al. 2019; Xiong et al. 2021; Wenke et al. 2022). We observed several described diazotroph bacteria in the sorghum mucilage, including *Acinetobacter*, *Curtobacterium*, *Herbaspirillum*, *Pantoea, Pseudomonas, Strenotrophomonas*, which were reported to colonize the maize mucilage (van Deynze et al. 2018; Higdon et al. 2020b, 2020a).

Regarding the fungal microbiome in the mucilage, we observed that the year of collection had the highest explanatory value. With two years of field data, there is not enough information to understand if the fungal community is responsive to other covariates (e.g., weather) or more stochastically assembled every year. Fungal community members likely have different responses than bacterial members to changing environmental conditions, including temperature, moisture, solar radiation, and precipitation (Jackson and Denney 2011; Copeland et al. 2015; Wagner et al. 2016; Gomes et al. 2018). We can deduce that the bacterial and fungal communities did not have strong relationships or co-dependencies based on their structures, and likely have different dominating drivers. However, the possibility of redundant functional relationships between different bacterial and fungal mucilage members cannot be eliminated.

We combined both culture-independent and dependent approaches to improve our understanding of the microbiome structure in phyllosphere exudates. Due to the chemical composition, plant DNA contamination, and low bacterial biomass associated with the wax and mucilage, a metagenomic sequencing approach would have been challenging to pursue with the sorghum phyllosphere (Sharpton 2014; van Deynze et al. 2018). Sequencing the V4 16S rRNA and the ITS1 regions allowed us to deeply characterize bacterial and fungal communities in sorghum phyllosphere exudates, albeit with limited taxonomic resolution that can be provided by the amplicons (to approximately the genus level Poretsky et al. 2014) as well as limited functional insight (Langille et al. 2013; Turner et al. 2013). Thus, we decided to culture wax and mucilage bacteria by using a variety of isolation media and growing conditions that we expected to enrich for plant-beneficial bacterial phenotypes. In the end, we were able to capture representatives of most of the bacterial taxa, at the family and genus levels, that we observed in our culture-independent approach. These isolates can now be used to test directly for plant beneficial properties and microbe-plant interactions in the laboratory, as well as to design synthetic communities to test as agricultural bioinoculants.

In summary, we report a characterization of microbiome structure of energy sorghum phyllosphere exudates, epicuticular wax and aerial root mucilage under multiple field conditions (irrigation, fertilization) and across two seasons for mucilage. We found that the wax and mucilage harbor distinct bacterial communities, suggesting niche specialization in the sorghum phyllosphere, and captured several key bacterial lineages in a parallel cultivation effort. Additionally, we found that fungal communities and bacterial communities in the mucilage are responsive to different drivers, with bacterial communities most distinctive by developmental stage and fungal communities most distinctive by year of sample collection. Next steps are to use the ecological dynamics from the cultivation-independent sequencing and apparent phenotypes of the bacterial isolates to understand the roles of these habitat-specialized microbiome members for plant performance and support the long-term goal of developing a carbon neutral/negative annual bioenergy crop.

## ACKNOWLEDGMENTS

This work was supported by the Great Lakes Bioenergy Research Center, U.S. Department of Energy, Office of Science, Office of Biological and Environmental Research under Award Number DE-SC0018409. Support for field research was provided by the Great Lakes Bioenergy Research Center, U.S. Department of Energy, Office of Science, Office of Biological and Environmental Research (Awards DE-SC0018409 and DE-FC02-07ER64494), by the National Science Foundation Long-term Ecological Research Program (DEB 1637653) at the Kellogg Biological Station, and by Michigan State University AgBioResearch. JM acknowledges field support from graduate students at TAMU. AS acknowledges support from Michigan State University AgBioResearch.

The authors declare no conflict of interest.

## SUPPLEMENTARY MATERIAL

**Supplementary Figure 1.** Relative abundance (RA) of microbial taxa at the genus level in sorghum epicuticular wax and aerial root mucilage. A) RA of Epicuticular wax bacterial communities. Samples were rarefied to 1,303 reads per sample. B) RA of aerial root mucilage bacterial communities. Samples were rarefied to 20,519 reads per sample. C) RA of aerial root mucilage fungal communities. Samples were rarefied to 33,975 reads per sample.

**Supplementary Table 1:** Pairwise comparisons using Wilcoxon rank sum test with continuity correction for differences in alpha diversity.

**Supplementary Table 2:** Media and conditions used in this study to culture the wax and mucilage microbiomes.

**Supplementary Table 3:** Relative abundance of bacterial taxa present in culture-independent and dependent approaches. Family level. Cut-off families < 0.1%

**Supplementary Table 4:** List of bacterial strains isolated from sorghum epicuticular wax and aerial root mucilage. Isolates were taxonomically identified by sequencing the full-length 16S rRNA gene.

## REFERENCES

Amicucci, M. J., Galermo, A. G., Guerrero, A., Treves, G., Nandita, E., Kailemia, M. J., et al. 2019. Strategy for Structural Elucidation of Polysaccharides: Elucidation of a Maize Mucilage that Harbors Diazotrophic Bacteria. Analytical Chemistry. 91:7254–7265 Available at: https://doi.org/10.1021/acs.analchem.9b00789.

Avato, P., Bianchi, G., and Mariani, G. 1984. Epicuticular waxes of Sorghum and some compositional changes with plant age. Phytochemistry. 23:2843–2846 Available at: https://www.sciencedirect.com/science/article/pii/0031942284830265.

Beattie, G. A., and Marcell, L. M. 2002. Effect of alterations in cuticular wax biosynthesis on the physicochemical properties and topography of maize leaf surfaces. Plant, Cell and Environment. 25:1–16.

Bennett, A. B., Pankievicz, V. C. S., and Ané, J.-M. 2020. A Model for Nitrogen Fixation in Cereal Crops. Trends in Plant Science. 25:226–235 Available at: https://www.sciencedirect.com/science/article/pii/S1360138519303292.

Bianchi, G., Avato, P., Bertorelli, P., and Mariani, G. 1978. Epicuticular waxes of two sorghum varieties. Phytochemistry. 17:999–1001 Available at: https://www.sciencedirect.com/science/article/pii/S0031942200886689.

Bolyen, E., Rideout, J. R., Dillon, M. R., Bokulich, N. A., Abnet, C. C., Al-Ghalith, G. A., et al. 2019. Reproducible, interactive, scalable and extensible microbiome data science using QIIME 2. Nature Biotechnology. 37:852–857 Available at: https://doi.org/10.1038/s41587-019-0209-9.

Bondada, B. R., Oosterhuis, D. M., Murphy, J. B., and Kim, K. S. 1996. Effect of water stress on the epicuticular wax composition and ultrastructure of cotton (Gossypium hirsutum L.) leaf, bract, and boll. Environmental and Experimental Botany. 36:61–69 Available at: https://www.sciencedirect.com/science/article/pii/0098847296001281.

Callahan, B. J., McMurdie, P. J., Rosen, M. J., Han, A. W., Johnson, A. J. A., and Holmes, S. P. 2016. DADA2: High-resolution sample inference from Illumina amplicon data. Nature Methods. 13:581–583 Available at: https://doi.org/10.1038/nmeth.3869.

Capella-Gutiérrez, S., Silla-Martínez, J. M., and Gabaldón, T. 2009. trimAl: a tool for automated alignment trimming in large-scale phylogenetic analyses. Bioinformatics. 25:1972–1973 Available at: https://doi.org/10.1093/bioinformatics/btp348.

Chai, Y. N., and Schachtman, D. P. 2022. Root exudates impact plant performance under abiotic stress. Trends in Plant Science. 27:80–91 Available at: https://doi.org/10.1016/j.tplants.2021.08.003.

Copeland, J. K., Yuan, L., Layeghifard, M., Wang, P. W., and Guttman, D. S. 2015. Seasonal Community Succession of the Phyllosphere Microbiome. Molecular Plant-Microbe Interactions®. 28:274–285 Available at: https://doi.org/10.1094/MPMI-10-14-0331-FI.

van Deynze, A., Zamora, P., Delaux, P.-M., Heitmann, C., Jayaraman, D., Rajasekar, S., et al. 2018. Nitrogen fixation in a landrace of maize is supported by a mucilage-associated diazotrophic microbiota. PLOS Biology. 16:e2006352–Available at: https://doi.org/10.1371/journal.pbio.2006352.

Dixon, P. 2003. VEGAN, A Package of R Functions for Community Ecology. Journal of Vegetation Science. 14:927–930 Available at: http://www.jstor.org/stable/3236992.

Doan, H. K., Ngassam, V. N., Gilmore, S. F., Tecon, R., Parikh, A. N., and Leveau, J. H. J. 2020. Topography-Driven Shape, Spread, and Retention of Leaf Surface Water Impacts Microbial Dispersion and Activity in the Phyllosphere. Phytobiomes Journal. 4:268–280 Available at: https://doi.org/10.1094/PBIOMES-01-20-0006-R.

Farber, C., Li, J., Hager, E., Chemelewski, R., Mullet, J., Rogachev, A. Yu., et al. 2019a. Complementarity of Raman and Infrared Spectroscopy for Structural Characterization of Plant Epicuticular Waxes. ACS Omega. 4:3700–3707 Available at: https://doi.org/10.1021/acsomega.8b03675.

Farber, C., Wang, R., Chemelewski, R., Mullet, J., and Kurouski, D. 2019b. Nanoscale Structural Organization of Plant Epicuticular Wax Probed by Atomic Force Microscope Infrared Spectroscopy. Analytical Chemistry. 91:2472–2479 Available at: https://doi.org/10.1021/acs.analchem.8b05294.

Galloway, A. F., Knox, P., and Krause, K. 2020. Sticky mucilages and exudates of plants: putative microenvironmental design elements with biotechnological value. New Phytologist. 225:1461–1469 Available at: https://doi.org/10.1111/nph.16144.

Gomes, T., Pereira, J. A., Benhadi, J., Lino-Neto, T., and Baptista, P. 2018. Endophytic and Epiphytic Phyllosphere Fungal Communities Are Shaped by Different Environmental Factors in a Mediterranean Ecosystem. Microbial Ecology. 76:668–679 Available at: https://doi.org/10.1007/s00248-018-1161-9.

Grady, K. L., Sorensen, J. W., Stopnisek, N., Guittar, J., and Shade, A. 2019. Assembly and seasonality of core phyllosphere microbiota on perennial biofuel crops. Nature Communications. 10:4135 Available at: https://doi.org/10.1038/s41467-019-11974-4.

Gregory, C. J., L, L. C., A, W. W., Donna, B.-L., A, L. C., J, T. P., et al. 2011. Global patterns of 16S rRNA diversity at a depth of millions of sequences per sample. Proceedings of the National Academy of Sciences. 108:4516–4522 Available at: https://doi.org/10.1073/pnas.1000080107.

Higdon, S. M., Pozzo, T., Kong, N., Huang, B. C., Yang, M. L., Jeannotte, R., et al. 2020a. Genomic characterization of a diazotrophic microbiota associated with maize aerial root mucilage. PLOS ONE. 15:e0239677–Available at: https://doi.org/10.1371/journal.pone.0239677.

Higdon, S. M., Pozzo, T., Tibbett, E. J., Chiu, C., Jeannotte, R., Weimer, B. C., et al. 2020b. Diazotrophic bacteria from maize exhibit multifaceted plant growth promotion traits in multiple hosts. PLOS ONE. 15:e0239081–Available at: https://doi.org/10.1371/journal.pone.0239081.

Hoang, D. T., Chernomor, O., von Haeseler, A., Minh, B. Q., and Vinh, L. S. 2018. UFBoot2: Improving the Ultrafast Bootstrap Approximation. Molecular Biology and Evolution. 35:518–522 Available at: https://doi.org/10.1093/molbev/msx281.

Hone, H., Mann, R., Yang, G., Kaur, J., Tannenbaum, I., Li, T., et al. 2021. Profiling, isolation and characterisation of beneficial microbes from the seed microbiomes of drought tolerant wheat. Scientific Reports. 11:11916 Available at: https://doi.org/10.1038/s41598-021-91351-8.

Igiehon, N. O., Babalola, O. O., and Aremu, B. R. 2019. Genomic insights into plant growth promoting rhizobia capable of enhancing soybean germination under drought stress. BMC Microbiology. 19:159 Available at: https://doi.org/10.1186/s12866-019-1536-1.

Jackson, C. R., and Denney, W. C. 2011. Annual and Seasonal Variation in the Phyllosphere Bacterial Community Associated with Leaves of the Southern Magnolia (Magnolia grandiflora). Microbial Ecology. 61:113–122 Available at: https://doi.org/10.1007/s00248-010-9742-2.

Jenks, M. A., Joly, R. J., Peters, P. J., Rich, P. J., Axtell, J. D., and Ashworth, E. N. 1994. Chemically Induced Cuticle Mutation Affecting Epidermal Conductance to Water Vapor and Disease Susceptibility in Sorghum bicolor (L.) Moench. Plant Physiol. 105:1239–1245 Available at: https://pubmed.ncbi.nlm.nih.gov/12232280.

Jenks, M. A., Rich, P. J., Rhodes, D., Ashworth, E. N., Axtell, J. D., and Ding, C.-K. 2000a. Leaf sheath cuticular waxes on bloomless and sparse-bloom mutants of Sorghum bicolor. Phytochemistry. 54:577–584 Available at: https://www.sciencedirect.com/science/article/pii/S0031942200001539.

Jenks, M. A., Rich, P. J., Rhodes, D., Ashworth, E. N., Axtell, J. D., and Ding, C.-K. 2000b. Leaf sheath cuticular waxes on bloomless and sparse-bloom mutants of Sorghum bicolor. Phytochemistry. 54:577–584 Available at: https://www.sciencedirect.com/science/article/pii/S0031942200001539.

Jenks, M. A., Tuttle, H. A., and Feldmann, K. A. 1996. Changes in epicuticular waxes on wildtype and eceriferum mutants in Arabidopsis during development. Phytochemistry. 42:29–34 Available at: https://www.sciencedirect.com/science/article/pii/0031942295008985.

Jones, J. G. 1969. Studies on Lipids of Soil Micro-organisms with Particular Reference to Hydrocarbons. Journal of General Microbiology. 59:145–152 Available at: https://www.microbiologyresearch.org/content/journal/micro/10.1099/00221287-59-2-145 [Accessed June 20, 2022].

Jordan, W. R., Shouse, P. J., Blum, A., Miller, F. R., and Monk, R. L. 1984. Environmental Physiology of Sorghum. II. Epicuticular Wax Load and Cuticular Transpiration1. Crop Science. 24:cropsci1984.0011183X002400060038x Available at: https://doi.org/10.2135/cropsci1984.0011183X002400060038x.

Kalyaanamoorthy, S., Minh, B. Q., Wong, T. K. F., von Haeseler, A., and Jermiin, L. S. 2017. ModelFinder: fast model selection for accurate phylogenetic estimates. Nat Methods. 14:587–589 Available at: https://pubmed.ncbi.nlm.nih.gov/28481363.

Katoh, K., Misawa, K., Kuma, K., and Miyata, T. 2002. MAFFT: a novel method for rapid multiple sequence alignment based on fast Fourier transform. Nucleic Acids Research. 30:3059–3066 Available at: https://doi.org/10.1093/nar/gkf436.

Kent, J., Hartman, M. D., Lee, D. K., and Hudiburg, T. 2020. Simulated Biomass Sorghum GHG Reduction Potential is Similar to Maize. Environmental Science & Technology. 54:12456–12466 Available at: https://doi.org/10.1021/acs.est.0c01676.

Ku, L. X., Sun, Z. H., Wang, C. L., Zhang, J., Zhao, R. F., Liu, H. Y., et al. 2012. QTL mapping and epistasis analysis of brace root traits in maize. Molecular Breeding. 30:697–708 Available at: https://doi.org/10.1007/s11032-011-9655-x.

Kunst, L., and Samuels, A. L. 2003. Biosynthesis and secretion of plant cuticular wax. Progress in Lipid Research. 42:51–80 Available at: https://www.sciencedirect.com/science/article/pii/S0163782702000450.

Langille, M. G. I., Zaneveld, J., Caporaso, J. G., McDonald, D., Knights, D., Reyes, J. A., et al. 2013. Predictive functional profiling of microbial communities using 16S rRNA marker gene sequences. Nature Biotechnology. 31:814–821 Available at: https://doi.org/10.1038/nbt.2676.

Letunic, I., and Bork, P. 2021. Interactive Tree Of Life (iTOL) v5: an online tool for phylogenetic tree display and annotation. Nucleic Acids Research. 49:W293–W296 Available at: https://doi.org/10.1093/nar/gkab301.

Levy, A., Salas Gonzalez, I., Mittelviefhaus, M., Clingenpeel, S., Herrera Paredes, S., Miao, J., et al. 2017. Genomic features of bacterial adaptation to plants. Nat Genet. 50:138–150 Available at: https://pubmed.ncbi.nlm.nih.gov/29255260.

Li, G., Li, L., Tarozo, R., Longo, W. M., Wang, K. J., Dong, H., et al. 2018. Microbial production of long-chain n-alkanes: Implication for interpreting sedimentary leaf wax signals. Organic Geochemistry. 115:24–31 Available at: https://www.sciencedirect.com/science/article/pii/S0146638017303819.

Li, W., and Godzik, A. 2006. Cd-hit: a fast program for clustering and comparing large sets of protein or nucleotide sequences. Bioinformatics. 22:1658–1659 Available at: https://doi.org/10.1093/bioinformatics/btl158.

Lindow, S. E., and Brandl, M. T. 2003. Microbiology of the phyllosphere. Appl Environ Microbiol. 69:1875–1883 Available at: https://pubmed.ncbi.nlm.nih.gov/12676659.

Love, M. I., Huber, W., and Anders, S. 2014. Moderated estimation of fold change and dispersion for RNA-seq data with DESeq2. Genome Biology. 15:550 Available at: https://doi.org/10.1186/s13059-014-0550-8.

Luo, Y., Wang, F., Huang, Y., Zhou, M., Gao, J., Yan, T., et al. 2019. Sphingomonas sp. Cra20 Increases Plant Growth Rate and Alters Rhizosphere Microbial Community Structure of Arabidopsis thaliana Under Drought Stress. Frontiers in Microbiology. 10 Available at: https://www.frontiersin.org/article/10.3389/fmicb.2019.01221.

Mahmoudi, T. R., Yu, J. M., Liu, S., Pierson, L. S., and Pierson, E. A. 2019. Drought-Stress Tolerance in Wheat Seedlings Conferred by Phenazine-Producing Rhizobacteria. Frontiers in Microbiology. 10 Available at: https://www.frontiersin.org/article/10.3389/fmicb.2019.01590.

McMurdie, P. J., and Holmes, S. 2013. phyloseq: An R Package for Reproducible Interactive Analysis and Graphics of Microbiome Census Data. PLOS ONE. 8:e61217–Available at: https://doi.org/10.1371/journal.pone.0061217.

Miller, C. S., Handley, K. M., Wrighton, K. C., Frischkorn, K. R., Thomas, B. C., and Banfield, J. F. 2013. Short-Read Assembly of Full-Length 16S Amplicons Reveals Bacterial Diversity in Subsurface Sediments. PLOS ONE. 8:e56018–Available at: https://doi.org/10.1371/journal.pone.0056018.

Minh, B. Q., Schmidt, H. A., Chernomor, O., Schrempf, D., Woodhams, M. D., von Haeseler, A., et al. 2020. IQ-TREE 2: New Models and Efficient Methods for Phylogenetic Inference in the Genomic Era. Molecular Biology and Evolution. 37:1530–1534 Available at: https://doi.org/10.1093/molbev/msaa015.

Mullet, J., Morishige, D., McCormick, R., Truong, S., Hilley, J., McKinley, B., et al. 2014. Energy Sorghum—a genetic model for the design of C4 grass bioenergy crops. Journal of Experimental Botany. 65:3479–3489 Available at: https://doi.org/10.1093/jxb/eru229.

Nazari, M., Riebeling, S., Banfield, C. C., Akale, A., Crosta, M., Mason-Jones, K., et al. 2020. Mucilage Polysaccharide Composition and Exudation in Maize From Contrasting Climatic Regions. Frontiers in Plant Science. 11 Available at: https://www.frontiersin.org/article/10.3389/fpls.2020.587610.

Nelle, V., Benjamin, C., Cheng, G., Grady, P., R, B. C., Dhruv, P., et al. 2019. Transcriptomic analysis of field-droughted sorghum from seedling to maturity reveals biotic and metabolic responses. Proceedings of the National Academy of Sciences. 116:27124–27132 Available at: https://doi.org/10.1073/pnas.1907500116.

Nilsson, R. H., Larsson, K.-H., Taylor, A. F. S., Bengtsson-Palme, J., Jeppesen, T. S., Schigel, D., et al. 2019. The UNITE database for molecular identification of fungi: handling dark taxa and parallel taxonomic classifications. Nucleic Acids Research. 47:D259–D264 Available at: https://doi.org/10.1093/nar/gky1022.

Olson, S. N., Ritter, K., Rooney, W., Kemanian, A., McCarl, B. A., Zhang, Y., et al. 2012. High biomass yield energy sorghum: developing a genetic model for C4 grass bioenergy crops. Biofuels, Bioproducts and Biorefining. 6:640–655 Available at: https://doi.org/10.1002/bbb.1357.

Peters, P. J., Jenks, M. A., Rich, P. J., Axtell, J. D., and Ejeta, G. 2009. Mutagenesis, Selection, and Allelic Analysis of Epicuticular Wax Mutants in Sorghum. Crop Science. 49:1250–1258 Available at: https://doi.org/10.2135/cropsci2008.08.0461.

Pierce, M. P. 2019. The ecological and evolutionary importance of nectar-secreting galls. Ecosphere. 10:e02670 Available at: https://doi.org/10.1002/ecs2.2670.

Poretsky, R., Rodriguez-R, L. M., Luo, C., Tsementzi, D., and Konstantinidis, K. T. 2014. Strengths and Limitations of 16S rRNA Gene Amplicon Sequencing in Revealing Temporal Microbial Community Dynamics. PLOS ONE. 9:e93827–Available at: https://doi.org/10.1371/journal.pone.0093827.

Punnuri, S., Harris-Shultz, K., Knoll, J., Ni, X., and Wang, H. 2017. The Genes Bm2 and Blmc that Affect Epicuticular Wax Deposition in Sorghum are Allelic. Crop Science. 57:1552–1556 Available at: https://doi.org/10.2135/cropsci2016.11.0937.

Quast, C., Pruesse, E., Yilmaz, P., Gerken, J., Schweer, T., Yarza, P., et al. 2013. The SILVA ribosomal RNA gene database project: improved data processing and web-based tools. Nucleic Acids Research. 41:D590–D596 Available at: https://doi.org/10.1093/nar/gks1219.

Reisberg, E. E., Hildebrandt, U., Riederer, M., and Hentschel, U. 2013a. Distinct phyllosphere bacterial communities on Arabidopsis wax mutant leaves. PLoS One. 8:e78613–e78613 Available at: https://pubmed.ncbi.nlm.nih.gov/24223831.

Reisberg, E. E., Hildebrandt, U., Riederer, M., and Hentschel, U. 2013b. Distinct phyllosphere bacterial communities on Arabidopsis wax mutant leaves. PLoS ONE. 8.

Reneau, J. W., Khangura, R. S., Stager, A., Erndwein, L., Weldekidan, T., Cook, D. D., et al. 2020. Maize brace roots provide stalk anchorage. Plant Direct. 4:e00284 Available at: https://doi.org/10.1002/pld3.284.

Rering, C. C., Beck, J. J., Hall, G. W., McCartney, M. M., and Vannette, R. L. 2018. Nectar-inhabiting microorganisms influence nectar volatile composition and attractiveness to a generalist pollinator. New Phytologist. 220:750–759 Available at: https://doi.org/10.1111/nph.14809.

Ruinen, J. 1965. The phyllosphere. Plant and Soil. 22:375–394 Available at: https://doi.org/10.1007/BF01422435.

Scully, M. J., Norris, G. A., Alarcon Falconi, T. M., and MacIntosh, D. L. 2021. Carbon intensity of corn ethanol in the United States: state of the science. Environmental Research Letters. 16:043001 Available at: http://dx.doi.org/10.1088/1748-9326/abde08.

Seemann, T. 2014. Prokka: rapid prokaryotic genome annotation. Bioinformatics. 30:2068–2069 Available at: https://doi.org/10.1093/bioinformatics/btu153.

Serrano, M., Coluccia, F., Torres, M., L’Haridon, F., and Métraux, J.-P. 2014. The cuticle and plant defense to pathogens. Front Plant Sci. 5:274 Available at: https://pubmed.ncbi.nlm.nih.gov/24982666.

Shanker, K. S., Kanjilal, S., Rao, B. V. S. K., Kishore, K. H., Misra, S., and Prasad, R. B. N. 2007. Isolation and antimicrobial evaluation of isomeric hydroxy ketones in leaf cuticular waxes of Annona squamosa. Phytochemical Analysis. 18:7–12 Available at: https://doi.org/10.1002/pca.942.

Sharpton, T. J. 2014. An introduction to the analysis of shotgun metagenomic data. Frontiers in Plant Science. 5 Available at: https://www.frontiersin.org/article/10.3389/fpls.2014.00209.

Shepherd, T., Robertson, G. W., Griffiths, D. W., Birch, A. N. E., and Duncan, G. 1995. Effects of environment on the composition of epicuticular wax from kale and swede. Phytochemistry. 40:407–417 Available at: https://www.sciencedirect.com/science/article/pii/003194229500281B.

Shepherd, T., and Wynne Griffiths, D. 2006. The effects of stress on plant cuticular waxes. New Phytologist. 171:469–499 Available at: https://doi.org/10.1111/j.1469-8137.2006.01826.x.

Stamp, P., and Kiel, C. 1992. Root Morphology of Maize and Its Relationship to Root Lodging. Journal of Agronomy and Crop Science. 168:113–118 Available at: https://doi.org/10.1111/j.1439-037X.1992.tb00987.x.

Steinmüller, D., and Tevini, M. 1985. Action of ultraviolet radiation (UV-B) upon cuticular waxes in some crop plants. Planta. 164:557–564 Available at: https://doi.org/10.1007/BF00395975.

Sun, A., Jiao, X.-Y., Chen, Q., Wu, A.-L., Zheng, Y., Lin, Y.-X., et al. 2021. Microbial communities in crop phyllosphere and root endosphere are more resistant than soil microbiota to fertilization. Soil Biology and Biochemistry. 153:108113 Available at: https://www.sciencedirect.com/science/article/pii/S0038071720304090.

Tsuba, M., Katagiri, C., Takeuchi, Y., Takada, Y., and Yamaoka, N. 2002. Chemical factors of the leaf surface involved in the morphogenesis of Blumeria graminis. Physiological and Molecular Plant Pathology. 60:51–57 Available at: https://www.sciencedirect.com/science/article/pii/S0885576502903760.

Turner, T. R., James, E. K., and Poole, P. S. 2013. The plant microbiome. Genome Biology. 14:209 Available at: https://doi.org/10.1186/gb-2013-14-6-209.

Ueda, H., Mitsuhara, I., Tabata, J., Kugimiya, S., Watanabe, T., Suzuki, K., et al. 2015. Extracellular esterases of phylloplane yeast Pseudozyma antarctica induce defect on cuticle layer structure and water-holding ability of plant leaves. Applied Microbiology and Biotechnology. 99:6405–6415 Available at: https://doi.org/10.1007/s00253-015-6523-3.

Vacher, C., Hampe, A., Porté, A. J., Sauer, U., Compant, S., and Morris, C. E. 2016a. The Phyllosphere: Microbial Jungle at the Plant–Climate Interface. Annual Review of Ecology, Evolution, and Systematics. 47:1–24 Available at: https://doi.org/10.1146/annurev-ecolsys-121415-032238.

Vacher, C., Hampe, A., Porté, A. J., Sauer, U., Compant, S., and Morris, C. E. 2016b. The Phyllosphere: Microbial Jungle at the Plant–Climate Interface. Annual Review of Ecology, Evolution, and Systematics. 47:1–24 Available at: https://doi.org/10.1146/annurev-ecolsys-121415-032238.

Vorholt, J. A. 2012. Microbial life in the phyllosphere. Nature Reviews Microbiology. 10:828–840 Available at: https://doi.org/10.1038/nrmicro2910.

Wagner, M. R., Lundberg, D. S., del Rio, T. G., Tringe, S. G., Dangl, J. L., and Mitchell-Olds, T. 2016. Host genotype and age shape the leaf and root microbiomes of a wild perennial plant. Nature Communications. 7:12151 Available at: https://doi.org/10.1038/ncomms12151.

Wang, X., Kong, L., Zhi, P., and Chang, C. 2020. Update on Cuticular Wax Biosynthesis and Its Roles in Plant Disease Resistance. Int J Mol Sci. 21:5514 Available at: https://pubmed.ncbi.nlm.nih.gov/32752176.

Weinstein, M. M., Prem, A., Jin, M., Tang, S., and Bhasin, J. M. 2019. FIGARO: An efficient and objective tool for optimizing microbiome rRNA gene trimming parameters. bioRxiv.:610394 Available at: http://biorxiv.org/content/early/2019/04/16/610394.abstract.

Wenke, S., Maria, S. L., Isabella, G., Karen, W., Marie, L., Babette, M., et al. 2022. Bacterial Succession and Community Dynamics of the Emerging Leaf Phyllosphere in Spring. Microbiology Spectrum. 10:e02420–21 Available at: https://doi.org/10.1128/spectrum.02420-21.

von Wettstein-Knowles, P. 1974. Ultrastructure and origin of epicuticular wax tubes. Journal of Ultrastructure Research. 46:483–498 Available at: https://www.sciencedirect.com/science/article/pii/S0022532074900690.

Xiong, C., Singh, B. K., He, J.-Z., Han, Y.-L., Li, P.-P., Wan, L.-H., et al. 2021. Plant developmental stage drives the differentiation in ecological role of the maize microbiome. Microbiome. 9:171 Available at: https://doi.org/10.1186/s40168-021-01118-6.

Xue, D., Zhang, X., Lu, X., Chen, G., and Chen, Z.-H. 2017. Molecular and Evolutionary Mechanisms of Cuticular Wax for Plant Drought Tolerance. Front Plant Sci. 8:621 Available at: https://pubmed.ncbi.nlm.nih.gov/28503179.

Yeats, T. H., and Rose, J. K. C. 2013. The formation and function of plant cuticles. Plant Physiol. 163:5–20 Available at: https://pubmed.ncbi.nlm.nih.gov/23893170.

