## Supplementary figures and images for "Phyllosphere exudates select for distinct microbiome members in sorghum epicuticular wax and aerial root mucilage"

### Supplementary Figure 1

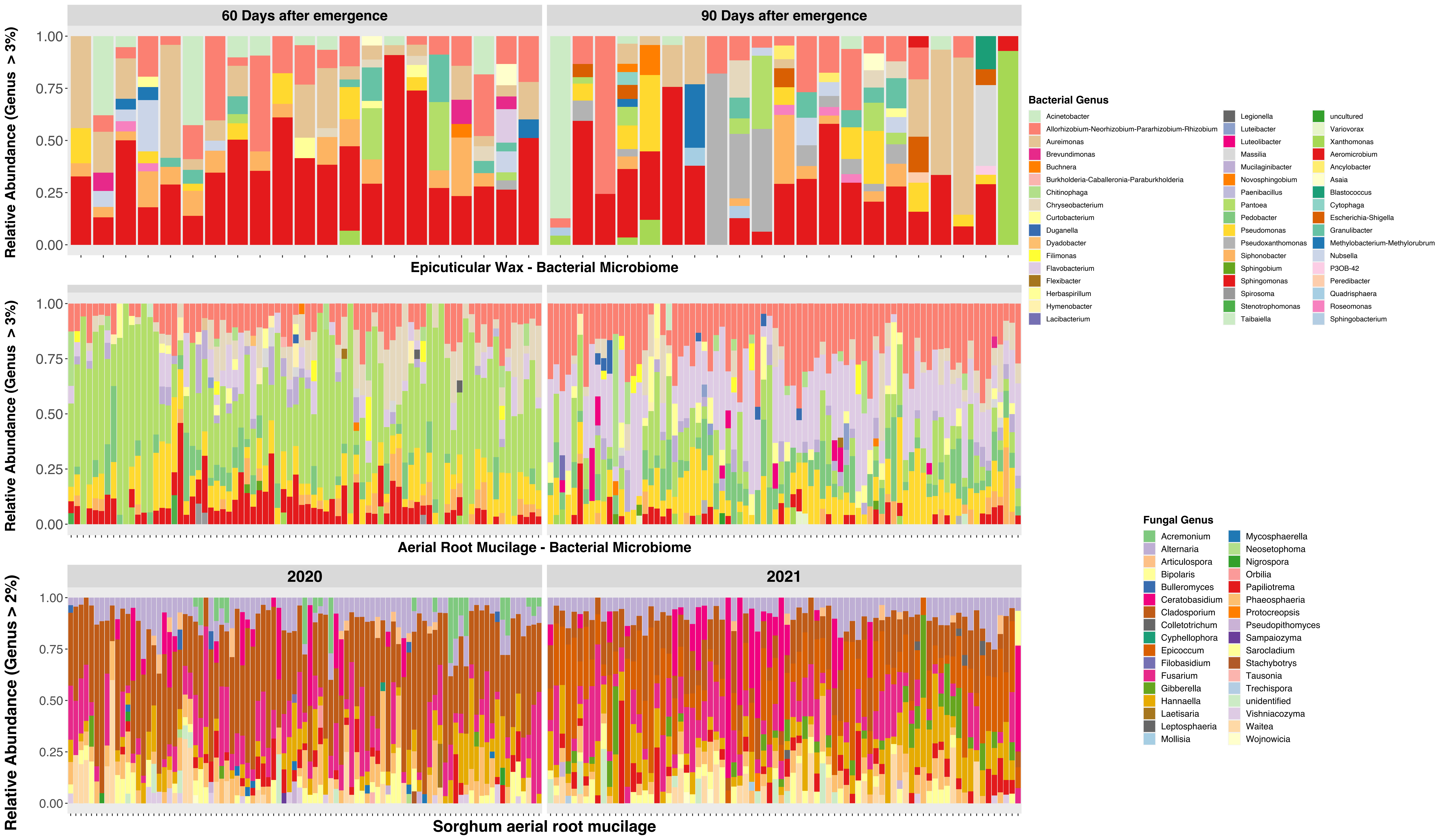
